# β-cell NCK1 is reduced in type 2 diabetes, impairing the unfolded protein response and insulin secretion while revealing sex-specific adaptation

**DOI:** 10.64898/2026.07.18.739339

**Authors:** L. Monteillet, R. Huet, J. Touzot, M. Schoumacher, L. Grieco-St-Pierre, C. Baldwin, C. Schmitt, M. Galipeau, E. Courty, S. Paraskevas, N. Jouvet, A. Güneş, J.L. Estall

## Abstract

Type 2 diabetes is characterized by failure of pancreatic β cells to adapt insulin secretion to metabolic demand, causing β cell function and reduced β cell mass. The unfolded protein response (UPR) is central to adaptation by maintaining endoplasmic reticulum homeostasis and supporting insulin biosynthesis, secretion, proliferation, and survival. NCK1, is an adaptor protein that regulates insulin biosynthesis and UPR activation, positioning it at the crossroads of several processes essential for β cell function. We found in mouse and human islets that NCK1 expression was upregulated by glucose and reduced in individuals living with type 2 diabetes. We thus explored the importance of β cell NCK1 in endocrine function and glucose homeostasis using INS-1 cells, mouse and human islets, and β cell–specific knockout male and female mice (NCK1βKO).

Loss of β cell NCK1 disrupted the UPR and impaired insulin gene expression, insulin content, and glucose-stimulated insulin secretion *in vitro*. Unexpectedly, these intrinsic β cell defects triggered sex-specific adaptive responses at the organismal level, revealing distinct strategies to reshape glucose homeostasis. Under high-fat/high-sucrose diet, both NCK1βKO female and male mice increased pancreatic insulin content, but females exhibited impaired insulin secretion associated with an increased number of small islets and reduced PERK pathway activation resulting in mild glucose intolerance. In contrast, males NCK1βKO mice had improved insulin secretion associated with islet expansion, β cell proliferation, and increased PERK pathway activation leading to enhanced glucose tolerance.

These findings identify NCK1 as a regulator of β cell insulin synthesis, secretion, and UPR signaling, and reveal sex-specific adaptive mechanisms to metabolic stress. Thus, reduced NCK1 in islets from people living type 2 diabetes may contribute to β cell dysfunction.

**RESEARCH IN CONTEXT:** *What is already known about this subject?:*

- β cell failure in type 2 diabetes results from impaired insulin secretion and inadequate adaptation of β cell mass to increased metabolic demand.
- The UPR, particularly the PERK pathway, is essential for maintaining β cell insulin biosynthesis, function, and adaptation to metabolic stress.
- NCK1 regulates protein translation, extracellular matrix re-organization and the UPR, and *in vitro* studies have suggested a possible role in β cell function.

*What is the key question?:*

- Does β cell NCK1 contribute to β cell metabolic stress responses and adaptive remodelling through its control of PERK signaling?

*What are the new findings?:*

- NCK1 expression is glucose-responsive and reduced in islets from individuals with type 2 diabetes.
- β cell NCK1 is required for efficient UPR activation, insulin biosynthesis, and glucose-stimulated insulin secretion.
- Loss of β cell NCK1 causes a sex-specific adaptive response to metabolic stress: males compensate through increased islet expansion, β cell proliferation, and enhanced insulin secretion; whereas females exhibit impaired insulin secretion despite preserved insulin stores. These differences are associated with sex-dependent regulation of PERK signaling and distinct patterns of islet remodeling.

*How might this impact clinical practice in the foreseeable future?:*

- Reduced β cell NCK1 can contribute to impaired β cell adaptation to metabolic stress, leading to hypoinsulinemia in type 2 diabetes. This pathway may represent a new mechanism of β cell failure and reveal novel targets to preserve insulin production and secretion in diabetes.

## INTRODUCTION

A hallmark of diabetes is failure of the pancreas to adapt insulin secretion to circulating glucose levels. This process relies on appropriate pancreatic β cell function and sufficient β cell mass. Defects in either of these can result in insufficient serum insulin, ultimately leading to diabetes^1^. One key biological pathway imperative for both β cell function and mass is the unfolded protein response (UPR)^2,3^, a signaling network that maintains endoplasmic reticulum (ER) homeostasis to ensure efficient insulin folding and processing. The UPR is activated when misfolded proteins accumulate and involves three pathways: IRE1α (type I transmembrane protein inositol requiring 1α), PERK (eukaryotic initiation factor 2α kinase), and ATF6 (activating transcription factor 6). Activation of these signaling arms reduces global protein synthesis, enhances expression of chaperones and folding enzymes, and promotes ER-associated degradation, restoring ER homeostasis. Experimental evidence demonstrates that an adaptive β cell UPR is essential, not only for preserving insulin production and glucose-stimulated insulin secretion but also for supporting proliferation and survival during periods of increased metabolic demand^4,5^. However, while the adaptive UPR response can be protective, chronic or excessive activation can trigger β cell failure and apoptosis^2,3^. Consequently, maladaptive ER stress is recognized as a major contributor to β cell failure and the progressive decline of β cell mass in diabetes^6,7^.

Several studies suggest that NCK1 (non-catalytic region of tyrosine kinase adaptor 1), an adaptor protein, plays an important role in β cell function and survival^8–10^. Through its scaffolding function, NCK1 assembles multi-protein complexes that control a wide range of cellular responses^11^, including actin cytoskeletal reorganization^12,13^, cell migration^14,15^, and proliferation^16,17^. NCK1 also has a critical role in mRNA translation^18–20^, the UPR^18,19,21^ and insulin signaling^22,23^. NCK1 interacts directly with PERK phosphorylated at Tyr-561, and *Nck1* silencing in murine beta cell lines enhances adaptive PERK activity, promotes proinsulin synthesis, and improves β cell survival under diabetogenic stress in cultured cells^9^. These data *in vitro* suggest that NCK1 may play an essential role in β cell health and function.

In this study, we investigated whether NCK1 expression is altered in type 2 diabetes and regulated by glucose using human islets, a publicly available cross-sectional single-cell RNA-sequencing dataset from human pancreatic islets, mouse islets, and INS-1 cells. We then investigated how the level of NCK1 affects UPR signaling in β cell insulin synthesis and secretion, and whole-body glucose homeostasis, using a β cell specific *Nck1* knockout mouse model (NCK1βKO).

## METHODS

### Mice

NCK1βKO mice were obtained by deleting *Nck1* exon 3 specifically in the pancreatic β cells. For that, C57BL/6N-A^tm1Brd^ Nck1^tm1a(EUCOMM)Wtsi^ KO first mice (MMRRC 48748) were cross-bred with the ACTB:FLPe B6J (Jackson laboratory 005703) to obtain C57BL/6JN Nck1^tm1c^ (Nck1^ex3fl/ex3fl^) mice. These were then crossed with the C57BL/6N-Ins1-Cre^Thor^ mice (Jackson Labs – 026801^24^) and backcrossed 10 times to C57BL/6N to obtain C57BL/6N Nck1^ex3fl/ex3fl^:Ins1-Cre^Thor^ mice, referred as NCK1βKO.

NCK1βKO mice were compared to age- and sex-matched Ins1-Cre^Thor^ (CRE) control mice from the same colony. Mice were housed in groups of 4-5 mice in an enriched environment, with free access to water, under controlled temperature conditions (21-23 °C) and with a 12h day/12 h night cycle. Littermates mice were randomly allocated to experimental groups and maintained on an *ad libitum* chow (Teklad Global 18% Protein - Rodent Diet) or high fat/high sucrose diet (D12451 – Rodent diet), as specified. Mice within each group were housed in multiple cages to minimize cage-related effects. Cages were maintained in the same animal room and on the same rack. The order of testing was randomized, irrespective of dietary group, to avoid order-related bias. Blinding was not performed during the study. Mice were monitored biweekly for body weight, general health, and signs of pain or distress throughout the study. Humane endpoints were defined according to IRCM animal care committee guidelines and included excessive weight loss or clinical signs of severe distress requiring euthanasia. No unexpected adverse events occurred, and no animals met the criteria for humane euthanasia. All experiments were performed in accordance with IRCM animal facility institutional animal care and use committee regulations (IRCM, 2022-03).

### Mouse primary islet isolation

Pancreata of 16–20-week-old mice were digested following perfusion with 0.4 U/ml Liberase TL (Millipore Sigma) in Hank’s Balanced Salt Solution (HBSS, Wisent) buffer (Ca2+/Mg2+) and dispersed by gentle shaking in buffer without Ca2+/Mg2+ containing 0.1% BSA and 20 mM HEPES, pH 7.4. Pellet was washed in HBSS buffer and resuspended in Histopaque-1077 (Sigma) overlaid with RPMI 1640 (without glucose) prior to separation by centrifugation. Handpicked islets were cultured overnight in 11 mM or 5 mM glucose RPMI 1640 (10% FBS and 1% penicillin/streptomycin) for recovery before experimentation. For each experiment, islets from one mouse represent one biological replicate. When an experiment required more islets than were obtainable from one mouse, islets from multiple mice were pooled to achieve one biological replicate.

### Static *in vitro* insulin secretion and insulin content

24 h after recovery time, batches of ten islets were equilibrated twice for 45 minutes in KRBH solution containing 0.1% (wt/vol.) BSA and 2.8 mmol/l glucose prior to incubation in 2.8 mmol/L glucose, 16.7 mmol/L glucose, or 16.7 mmol/L glucose with 25 mmol/L KCl at 37 °C for 1 h. Intracellular insulin content was measured after lysis in acid–ethanol (1.5% HCl in 70% EtOH) using the mouse insulin ELISA immunoassay kit (Alpco or Promega). For pancreatic insulin, half of the pancreas was incubated overnight at −20°C in 5 mL acid-ethanol (1.5% HCl in 70% EtOH) prior to homogenization and refreezing. Thawed homogenate was centrifuged prior to neutralization with 1 mol/L Tris pH 7.5. Insulin was measured by immunoassay ELISA kit (Promega) and normalized to pancreas weight.

### Cell treatments

All INS-1 cell lines^25^ were cultured at 37 °C with 5% CO2 in RPMI-1640 supplemented with 2 mM L-Glutamine (Wisent), 1 mM sodium pyruvate, 10 mM HEPES, 0.05 mM β-mercaptoethanol, 10% HI-FBS, and 1% Pen/Strep. INS-1 cells or mouse islets were treated for indicated times with: 1 μM thapsigargin (Sigma, T9033) or its vehicle control 70% ethanol for 24 h; different concentrations of salubrinal (Sigma, SML0951), as indicated, or its vehicle DMSO for 18 h (BioShop, DMS666.100); 2 µM of PERK inhibitor GSK2656157 (Cayan medical, 17372) or its vehicle DMSO; or, with palmitate (1 mM or 0.5 mM) (Sigma, P9767) or its vehicle control bovine serum albumin (BSA; Sigma-Aldrich) with or without high glucose (30 mM). All INS-1 cell lines were routinely tested for mycoplasma contamination at least once per year and were confirmed to be mycoplasma-free throughout the study.

### INS-1 NCK1 KD cell lines

Stable or acute INS-1 NCK1-KD cell lines were obtained by infecting, at an MOI of 5, the parental INS-1 cell line^25^ with lentiviruses containing a control shRNA or different shRNA against *Nck1* mRNA **(Table 3)**. For the stable INS-1 NCK1-KO cell lines, cells were selected with puromycin (0.5 μg/mL).

### Protein isolation and western blotting

Total cell lysates were prepared using RIPA Buffer supplemented with protease and phosphatase inhibitors, diluted in Laemmli sample buffer containing 10% β-mercaptoethanol and denatured at 95 °C for 10 minutes. For non-reducing gels, (misfolded proinsulin), cells or islets were lysed directly in 1X Laemmli sample buffer without β-mercaptoethanol and denatured at 95 °C for 10 minutes. Proteins were separated by SDS-PAGE, transferred to 0.2 µM nitrocellulose membranes (Bio-Rad), blocked in TBS with 5% milk (BioShop) for a minimum of one hour, incubated with primary antibodies overnight and then 1 h with secondary antibodies (Supplementary **Table 1**). For islets, total proinsulin signal was normalized to total protein level, assessed by the Revert™ 700 Total Protein Stain (Li-cor, 926-11021), according to Li-cor protocol.

### Gene expression analysis

RNA was extracted from INS-1 cells using Trizol (Invitrogen) and from islets (∼120) by RNeasy Mini Kit (Qiagen) or acid guanidinium thiocyanate–phenol–chloroform extraction, as previously described^26^. cDNA was synthetize (High-capacity reverse transcription kit, Applied Biosystems) using 1 µg of RNA treated with DNAse-I (Thermo Fischer), quantified using Syber Green PCR mastermix (ABM, G891, G892) and a 384-well QuantStudio 5 (Life Technologies, Thermo Fischer) and presented as "relative mRNA expression" using the ΔΔCt threshold cycle method, with normalization to the Hypoxanthine-guanine phosphoribosyltransferase (*Hprt*) reference gene. Primers used for the qPCR data are listed in Supplementary **Table 2**.

### MTT viability assay

INS-1 cells were cultured for at least 24h before the start of the experiment. After treatments, cells incubated for 3 h at 37°C with a thiazolyl Blue reagent (MTT) (BioShop, MTT222.250) solution (prepared at 5 mg/ml in PBS) diluted 1:10 in regular medium (final concentration: 0.5 mg/mL). After two times with PBS the solubilization solution (40% dimethylformamide, 2% glacial acetic acid, 16% sodium dodecyl sulfate, pH 4.7) was added and absorbance was read at 570 nm. For each cell line, cell numbers were normalized to their own basal condition (5.5 mM glucose - DMSO). Data were expressed as a percentage of viable cells relative to control cells under glucolipotoxic condition.

### Metabolic tests and serum insulin

Oral glucose tolerance tests (oGTT), mixed meal tolerance test (MMTT) and insulin sensitivity tests were performed after a 6 h fast following oral administration of glucose (1.5 g/kg or 1g/kg for long term diet protocol) or Ensure® Plus Calories (7 μL/g = 1.4 g glucose/kg; 0.35 g protein/kg; 0.30 g lipid/kg), or injection of human insulin (0.7 unit/kg i.p. for males and 0.5 units/kg for females) respectively. Blood glucose and serum insulin were measured by tail vein using a standard glucometer (FreeStyle Lite, Abbott Diabetes Care) or the mouse ultrasensitive insulin ELISA (Alpco).

### Islet Area/Mass and Immunohistochemistry

For islet area, 10 sections of 5 μm of mouse pancreas (paraffin embedded), separated by at least 50 μm, were stained with hematoxylin-eosin. Islet density (number of islet/mm^2^) was calculated as the average number of islets per section divided by the section area. Islet area (%) was calculated as: 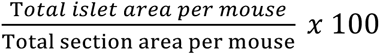

### Human islet culture

Human islets for research were provided by the Alberta Diabetes Institute IsletCore at the University of Alberta in Edmonton (http://www.bcell.org/adi-isletcore.html) with the assistance of the Human Organ Procurement and Exchange (HOPE) program, Trillium Gift of Life Network (TGLN), and other Canadian organ procurement organizations (Supplementary **Table 3-Human islets checklist**). Islet isolation from donor pancreata was approved by the Human Research Ethics Board at the University of Alberta (Pro00013094). All donors’ families gave informed consent for the use of pancreatic tissue in research. Human islets received from ADI Isletcore were allowed to recover in DMEM containing 5.5 mM glucose and supplemented with 10% FBS, 1% Pen/Strep overnight. To assess the level of NCK1 in response to glucose, human islets from one donor were picked, dispersed with accutase, plated on uncoated 12 well plates (∼150 IEQ/well) and incubated with DMEM containing 5.5 mM; 11 mM or 25 mM glucose in duplicate for 24 h.

### Human β cell scRNA-sequencing data

Single-cell gene expression correlations were analyzed using a publicly available dataset (GSE221156), consisting of single-cell transcriptome of 48 human pancreatic islets coming from 17 Healthy, 14 pre-diabetic and 17 type 2 diabetic cadaveric donors matched by sex, age and ancestry^27^. We had access only to released data and associated metadata. It is important to note that the original dataset was designed to characterize transcriptional changes associated with T2D and not to investigate gene-gene relationships examined in the present study. Consequently, relevant biological or technical variables may not have been measured or available for adjustment. Moreover, correlations derived from transcriptomic data do not imply causality, as they may reflect indirect regulatory relationships or shared responses to common biological processes. Therefore, the observed associations should be interpreted as hypothesis-generating. In addition, the large number of cells analyzed may yield statistically significant associations with limited biological relevance; therefore, Spearman’s correlation coefficient (ρ) and the corresponding pseudo-R² (ρ²) should be interpreted alongside p values.

Raw data and the original R-based processing pipeline were obtained from the Gene Expression Omnibus (GSE221156) and Zenodo (https://zenodo.org/records/14656366). Gene–gene correlations were assessed in normalized human β cell data using Spearman’s rank correlation coefficient and the cor.test function in R (version 4.4.2). Correlations were calculated both at the single-cell level and using the pseudo-bulk dataset processed as described by the original study. For each gene pair, Spearman’s ρ and the corresponding p value were calculated; ρ² was used as an approximate measure of explained variance.

### Statistical analysis

Individual data points identified as statistical outliers using Grubbs’ test (significance level α = 0.05, GraphPad Prism) were excluded from analyses. This exclusion criterion was established *a priori*. Mice that failed to gain weight due to dental problems were also excluded from analyses. This exclusion criterion was also established *a priori*. No other animal exclusion criteria were applied. To compare two groups, an unpaired *t*-test was applied. For the analysis of more than one gene or protein expression between two groups, multiple unpaired *t*-tests were conducted. When comparing more than two experimental groups with one variable, one-way ANOVA were employed. For experiments involving two or more groups and two variables, a two-way ANOVA was performed, followed by appropriate multiple comparisons tests. Post hoc analyses included two-tailed *t*-tests, Tukey’s test (for parametric one-way ANOVA test) and Tukey’s or Sidak’s multiple comparisons tests (for two-way ANOVA), as indicated.

## RESULTS

### β cell NCK1 increases in response to high glucose but is low in human islets from people living with type 2 diabetes

To investigate whether β cell NCK1 expression is altered in diabetes, we measured NCK1 levels in human islets. NCK1 protein levels were decreased in both female and male human islets from type 2 diabetic donors compared to healthy donors **(FIGURE 1A)**. Consistently, analysis of a published scRNAseq database^27^ with single-cell transcriptomes of human islets revealed that *NCK1* mRNA trended down in β cells from type 2 diabetic donors compared to healthy (*p-value = 0.15)* and prediabetic donors (*p-value = 0.12).* No such trend was observed in other endocrine cell populations **(FIGURE 1B, FIGURE S1A)**.

**Figure 1.**
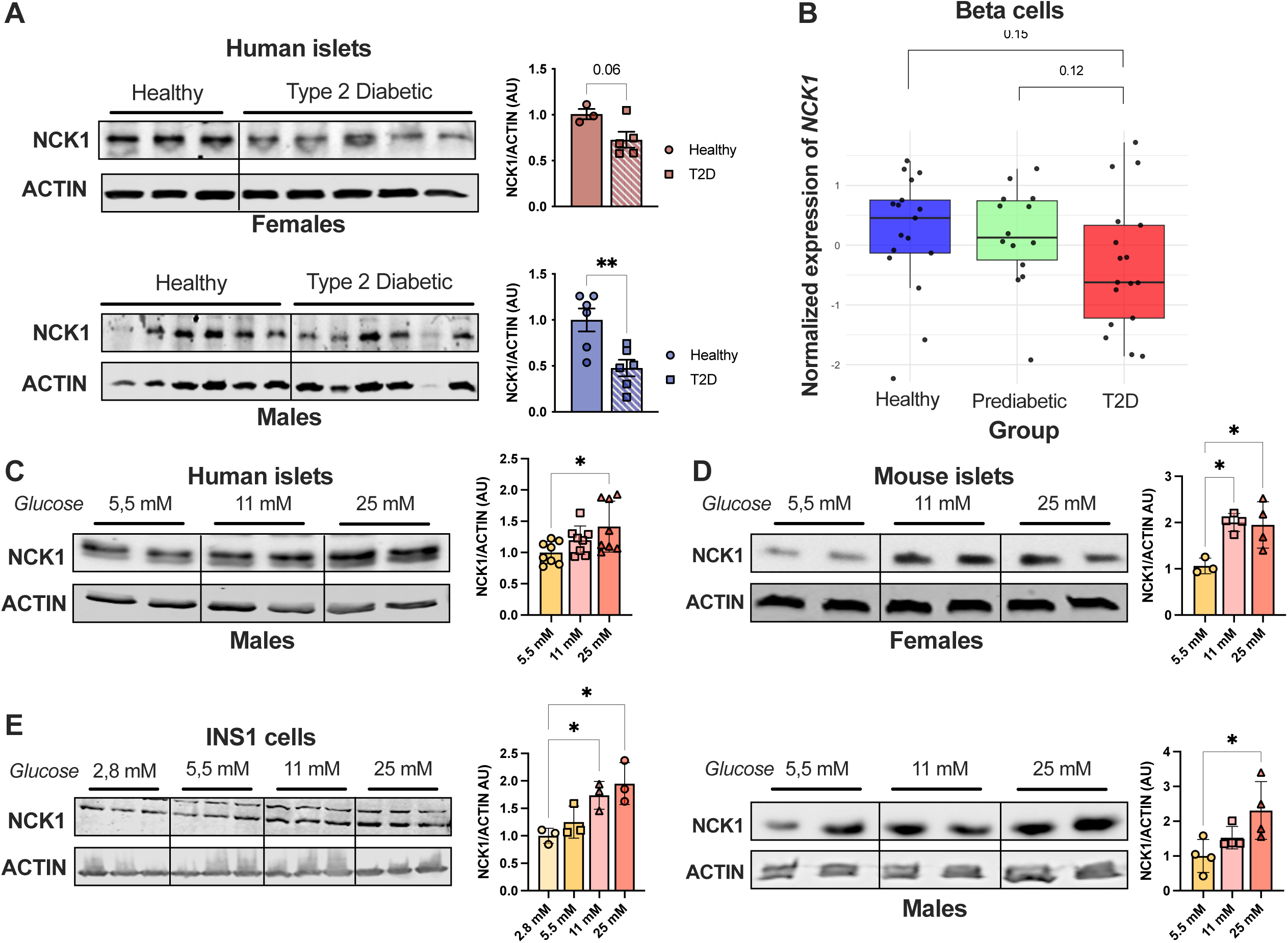
NCK1 expression is responsive to glucose in β cells and decreased in islets from T2D donors. **(A)** Relative protein expression of NCK1 in female and male human islets from donors healthy or people living with type 2 diabetes (Females, healthy n=3, T2D n=5; Males, heathy n=6, T2D n=6). **(B)** Pseudobulk analysis of *NCK1* expression in human β cells from healthy (n=17), prediabetic (n=14) and T2D (n=17) donors. **(C)** Relative protein expression of NCK1 in human islets from male healthy donors treated 24 h with glucose (5,5 mM; 11 mM or 25 mM) (n=3). Western blot images are representative of 3 independent experiments performed in duplicate. Bar charts are quantification of 3 independent experiments. **(D)** Relative protein expression of NCK1 in islets from female and male mice treated 48 h with glucose (5,5 mM; 11 mM aor 25 mM) (mixture of islets from 8 mice pooled together – repeated twice, n=2). Western blot images are representative of 2 independent experiments performed in duplicate. Bar charts are quantitation of 2 experiments. **(E)** Relative expression of NCK1 in INS-1 cells treated 48 h with glucose (2,8 mM; 5,5 mM; 11 mM or 25 mM) (n=3). Western blot images and bar charts are representative of 3 independent experiments performed in triplicate. ACTIN was used as the loading control to normalize data. Data are expressed as means ± SEM (A, B) or ± SD (C, D, E). Statistical significance was assessed using *t*-tests and one-way ANOVA, with *p ≤ 0.05 and **p ≤ 0.01.

Given previous studies implicating NCK1 as an important regulator of insulin processing^12,13^, decreased levels in diabetes prompted us to investigate mechanisms that regulate β cell NCK1 expression. In response to high glucose, NCK1 protein expression trended upward in healthy male human islets (**FIGURE 1C)** and was significantly increased in female and male mouse islets (**FIGURE 1D & S1B**) as well as in the rat INS-1 cell line (**FIGURE 1E & S1C**). Together, our findings indicated that NCK1 is positively regulated by glucose and reveal an association between low NCK1 and β cell dysfunction.

### Loss of NCK1 decreases insulin synthesis and secretion

We next assessed whether low NCK1 affects the capacity of β cells to synthetize insulin. In contrast to published work^8,10^, chronic reduction of *Nck1* in INS-1 cells caused decreases in insulin (**FIGURE 2A)** and proinsulin content (**FIGURE 2B),** as well as reduced mRNA of insulin genes (*Ins1, Ins2*) (**FIGURE 2C**), correlating with the efficiency of *Nck1* knockdown. Similar results were obtained following acute *Nck1* knockdown (**FIGURE S2**).

**Figure 2.**
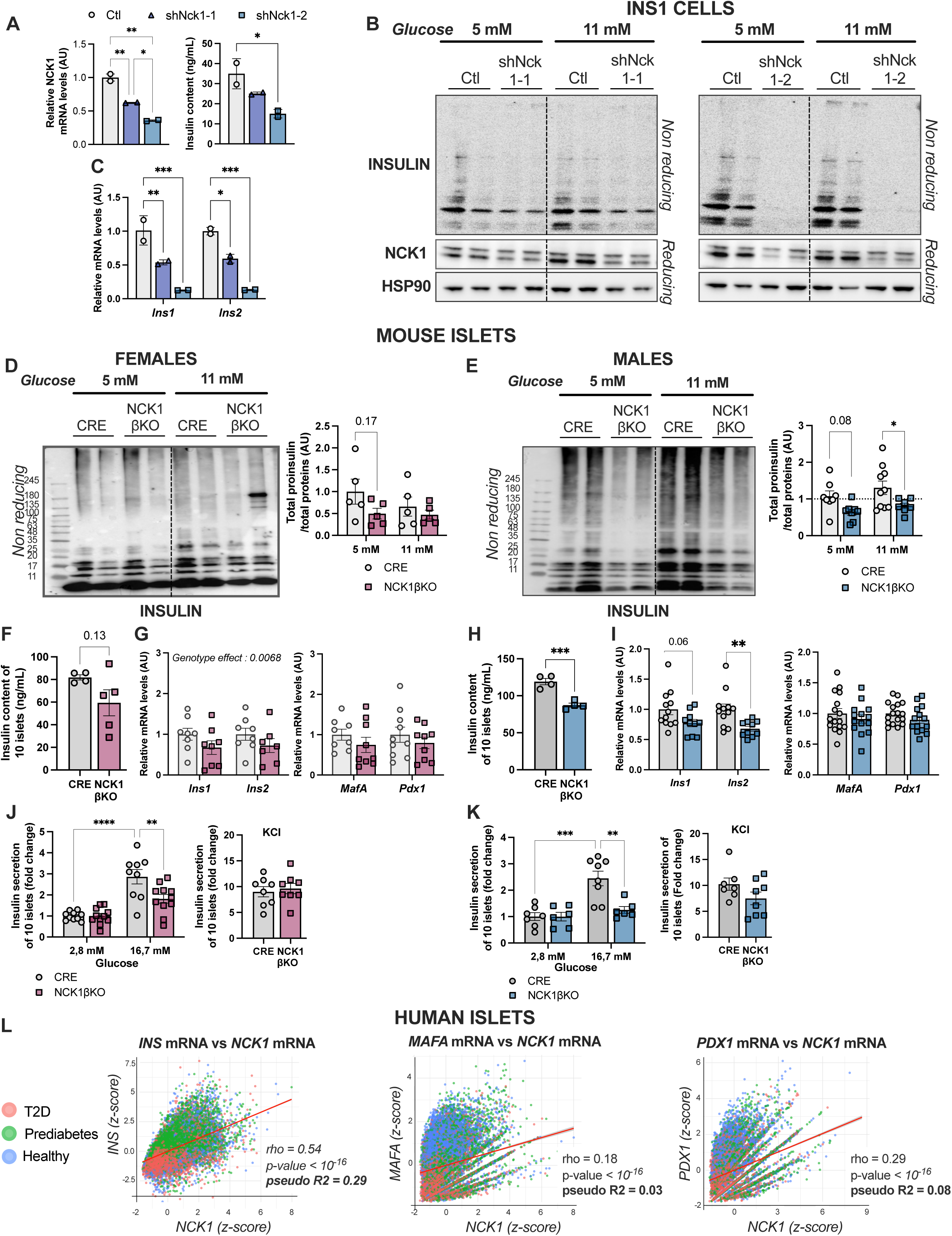
NCK1 deficiency impairs insulin production, proinsulin folding and β cell function. **(A)** Relative mRNA expression of *Nck1* and insulin content of stable NCK1-KD or control INS-1 cells in 5,5 mM glucose (n=4-5). **(B)** Representative images of proinsulin and insulin assessed by nonreducing SDS-PAGE of total cell lysate and NCK1 protein in stable NCK1-KD or control INS-1 cells incubated 8 h in 5 mM or 11 mM glucose (n=5). **(C)** Relative mRNA expression of insulin genes (*Ins1, Ins2*) of stable NCK1-KD or control INS-1 cells under 5,5 mM glucose condition measured by RT-qPCR (n=4-5). **(D, E)** Representative images and quantification of proinsulin and insulin assessed by nonreducing SDS-PAGE of total cell lysate from female and male NCK1βKO and CRE control islets incubated 24 h in 5 mM or 11 mM glucose (Females n=5; Males n=8-10). **(F, H)** Representative islet insulin content of 10 primary islets from female and male NCK1βKO or CRE control mice (Females n=4-5; Males n=4). **(G, I)** Relative mRNA expression of insulin genes (*Ins1, Ins2*) and insulin gene transcription factors (*Mafa*, *Pdx1*) of islets from female and male NCK1βKO or CRE control mice incubated 48 h in 5,5 mM glucose (Females n=7-10; Males n=10-17). **(J-K)** *In vitro* glucose-stimulated insulin secretion (GSIS) (Females n=10-11; Males n=6-8) in response to 2,8 mM or 16,7 mM of glucose or 16,7 mM of glucose + 25 mM KCl. **(L)** Correlative mRNA expression analysis of scRNAseq data from human β cells from 17 healthy, 14 pre-diabetic and 17 T2D cadaveric donor islets matched by sex, age and ancestry. Data are expressed as means ± SD (INS-1 cells) or SEM (islets). *Hprt* was used as an internal control to normalize qPCR data. Total insulin on the blots was normalized to protein content. Statistical significance was assessed using two-way ANOVA followed by a Tukey’s multiple comparisons test for GSIS, multiple t-tests for mouse islet data, and one-way ANOVA for INS-1 data, with *P ≤ 0.05, **P ≤ 0.01, ***P ≤ 0.001 and ****P ≤ 0.0001.

To determine how reduced NCK1 affects β cell endocrine function and whole-body glucose homeostasis, we created a mouse model where NCK1 was specifically and constitutively deleted in pancreatic β cells (NCK1βKO mice). Consistent with cell line data, loss of β cell NCK1 reduced total islet proinsulin and insulin content in female and male islets (**FIGURE 2D-F, H**). The expression of insulin genes (*Ins1, Ins2*) were also reduced (**FIGURE 2G, I**) and both female and male NCK1βKO islets had impaired insulin secretion in response to high glucose (16.7 mM) *ex vivo* (**FIGURE 2J-K**). In line with mouse data, human islet scRNAseq data revealed a positive association of *NCK1* with *INS* mRNA in β cells (**FIGURE 2L**). Together, these results indicate that NCK1 is a glucose-responsive protein required for the maintenance of β cell insulin expression, content and glucose-stimulated insulin secretion (GSIS).

### Loss of NCK1 in β cells impairs the unfolded protein response

NCK1 is characterized as a regulator of the UPR and directly interacts with PERK^8^. Since tight regulation of PERK, and the UPR in general, is crucial for insulin expression and secretion^2,28^, we assessed whether these pathways were affected in our models. In contrast to data in MIN6 cells^8,9^, reduction of NCK1 in INS-1 cells (stable or acute) or islets from female and male mice reduced basal PERK pathway activity, including PERK/eIF2α phosphorylation and expression of PERK target genes (*Atf4*, *Chop*, and *Gadd34*) (**FIGURE 3A-B; FIGURE 3 E-F & FIGURE S3A-B**). The IRE1α target gene *Hrd1* expression was also reduced across models, while additional UPR components, including *Atf6* and chaperone expression (*Bip* and *Grp94*), displayed model-dependent changes (**FIGURE 3C-D; FIGURE 3 G-H & FIGURE S3C**). This was in line with positive associations between *Nck1* and *Gadd34*, *Atf3, Chop* and *Hrd1* mRNA levels in scRNAseq data from human β cells (**FIGURE 3I**). Overall, these results suggest that the decrease in insulin expression and GSIS caused by low β cell NCK1 may be associated with reduced PERK activity.

**Figure 3.**
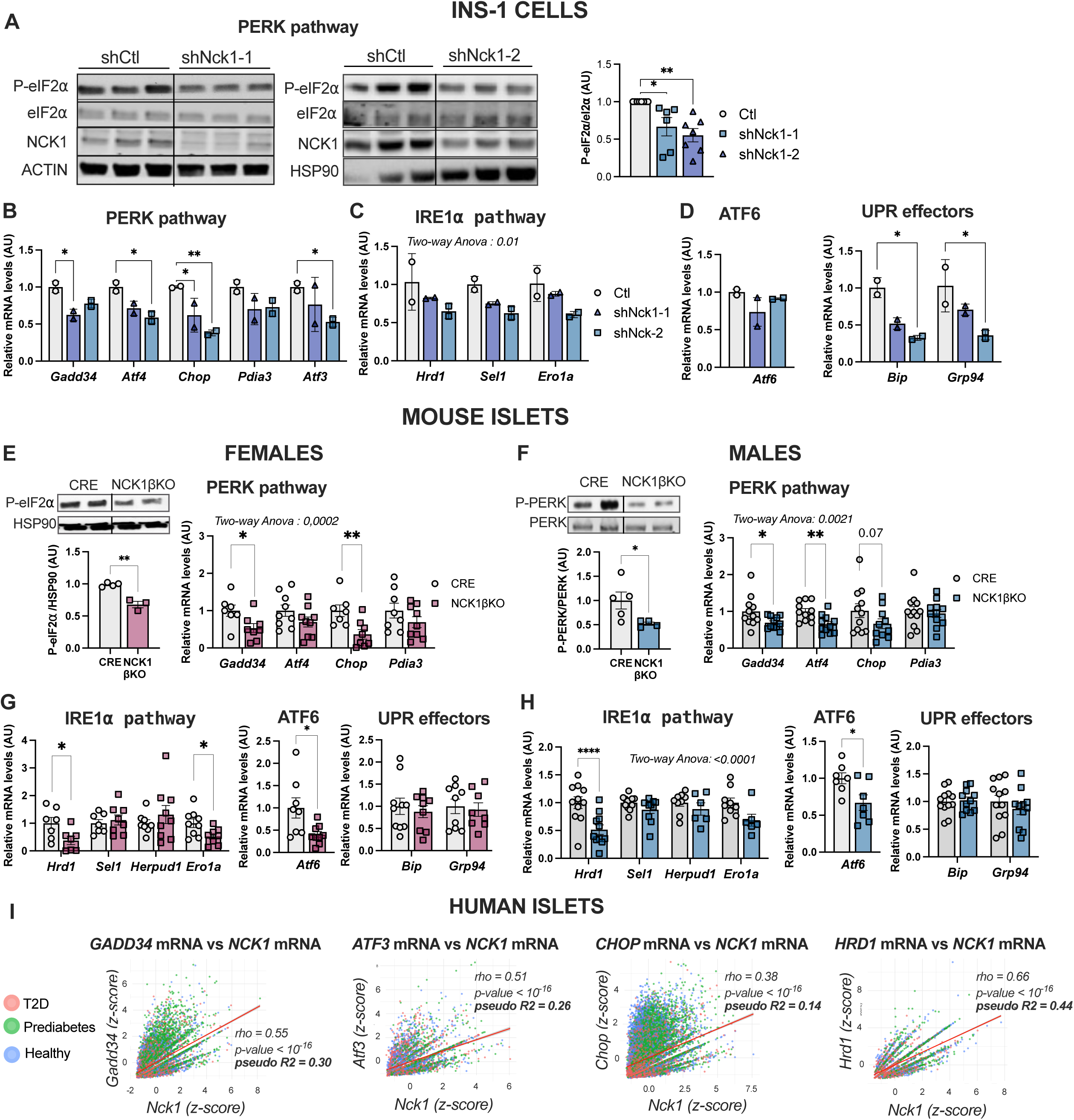
NCK1 deficiency downregulates the unfolded protein response in β cells. **(A)** PERK pathway activation measured by the ratio P-eIF2*a*/eIF2*a* and **(B)** PERK target genes expression in stable NCK1-KD or control INS-1 cells incubated 36 h in 5,5 mM of glucose. Western blots are representative of 6-7 independent experiments and bar charts are quantification 6-7 experiments. **(C)** IRE1 target mRNA expression, **(D)** *Atf6* and chaperone protein mRNA expression in stable NCK1-KD or control INS-1 cells incubated 36 h in 5,5 mM of glucose (n=5). **(E-F)** PERK pathway activation measured by the ratio P-eIF2*a*/HSP90 or P-PERK/PERK (females n=3-4; males n=4-5), target gene expression (females n=7-10; males n=10-12), **(G-H)** and IRE1 pathway target genes, *Atf6* and chaperone protein mRNA expression in isolated islets from female and male NCK1βKO and CRE control mice incubated 48 h in 5,5 mM of glucose (females n=7-10; males n=7-12). **(I)** Correlative mRNA expression analysis of scRNAseq data from human β cells of 17 healthy, 14 pre-diabetic and 17 T2D cadaveric donors islets matched by sex, age and ancestry. Data are expressed as means ± SD (INS-1) or SEM (islets). HSP90 was used as a loading control to normalize western blot data. *Hprt* was used as an internal control for qPCR. Statistical significance was assessed using one-way ANOVA for panels A and D and two-way ANOVA for other INS-1 cell data, followed by Tukey’s multiple comparison. Multiple t-tests were performed for the mouse islet data, with *p ≤ 0.05, **p ≤ 0.01, ***p ≤ 0.001 and ****p ≤ 0.0001.

### Sex influences how β cell NCK1 loss affects islet insulin content and secretion *in vivo*

We next examined whether the intrinsic β cell secretory defect caused by NCK1 deletion altered whole-body glucose homeostasis. When fed a chow diet, female and male NCK1βKO mice had body weights and fasting metabolic parameters comparable to CRE control mice (**FIGURE S4A–F**). Despite decreased GSIS in NCK1βKO primary islets from both sexes (**FIGURE 2J, K**), *in vivo* GSIS was sexually dimorphic. Consistent with the *ex vivo* islet phenotype, females NCK1βKO mice had reduced insulin secretion during a glucose challenge. However, this was accompanied by modest improvements in glucose tolerance (**FIGURE S4 G-H**) possibly due to increased insulin sensitivity (**FIGURE S4K**). In contrast, male NCK1βKO mice maintained normal *in vivo* insulin secretion and glucose homeostasis (**FIGURE S4 I-J, M**) together with increased pancreatic insulin content (**FIGURE S4N**), an adaptation not observed in females (**FIGURE S4L**).

Metabolic challenges (such as high caloric diets) place a large demand on β cells to sustain insulin production. Therefore, to determine whether β cell NCK1 loss impacts glucose homeostasis in conditions of increased insulin demand, mice were challenged with a short-term high fat/high sucrose (HF/HS) diet (**FIGURE 4A**). As expected, the HF/HS diet increased body weight (**FIGURE 4B, C**) and fasting glycemia (**FIGURE 4D, F**). The sex-specific phenotype observed under chow diet persisted. Female NCK1βKO mice had the same fasting insulinemia (**FIGURE 4E**) as controls but secreted less insulin during both a glucose (oGTT) or mixed meal (MMTT) challenge, resulting in a mild impairment of tolerance to glucose (**FIGURE 4H-K**). This was again within a setting of improved insulin sensitivity (**FIGURE 4L**). In contrast, male NCK1βKO mice had increased fasting insulin **(FIGURE 4G**) and normal glucose tolerance during both the oGTT and MMTT, with enhanced insulin secretion during MMTT (**FIGURE 4N-P**). Insulin sensitivity was unaffected by diet or genotype in males **(FIGURE 4Q**).

**Figure 4.**
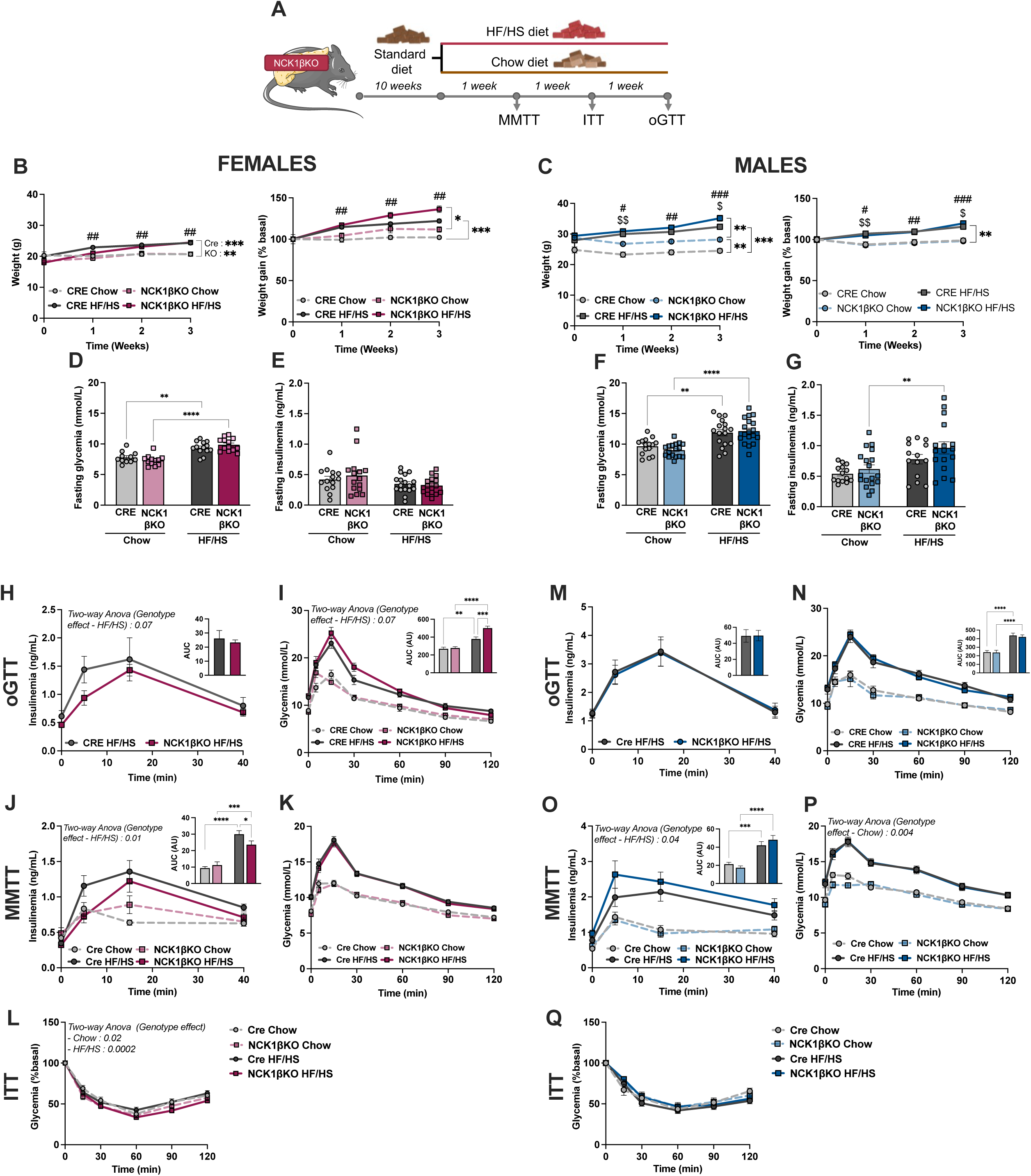
β cell NCK1 deficiency differentially regulates insulin secretion and glucose homeostasis in male and female mice during high-fat/high-sucrose feeding. **(A**) Time course of metabolic tests for male and female NCK1βKO and CRE control mice fed a chow or high-fat, high-sucrose (HF/HS) diet. **(B, C)** Body weight and weight gain, **(D, E)** 6 h-fasting glycemia and 6 h-fasting insulinemia of female and **(F, G)** male NCK1βKO and CRE control mice, after a 1-week chow or HF/HS diet (Females, n=14-18; Males, n= 15-18). **(H, M)** Insulinemia and **(I, N)** glycemia during an oral glucose tolerance test in female and male NCK1βKO and CRE control after 3 weeks of chow or HF/HS diet (Females, n=14-18; Males, n= 15-18). **(J, O)** Insulinemia and **(K,P)** glycemia during an oral mixed meal tolerance test on female and male NCK1βKO and CRE control mice, after a 1-week chow or HF/HS diet (Males, n= 7-8; Females, n=7-9). **(L, R)** Insulin tolerance in female and male NCK1βKO and CRE control mice, after 2 weeks of chow or HF/HS diet (Females, n=7-9; Males, n= 7-8). Data are expressed as means ± SEM. Statistical significance was assessed using two-way ANOVA followed by Tukey’s multiple comparison with *p ≤ 0.05, **p ≤ 0.01, ***p ≤ 0.001 and ****p ≤ 0.0001.

Mice were maintained on the HF/HS diet for an additional 19 weeks (**FIGURE 5A**) and the sexual dimorphism became more pronounced. Female NCK1βKO mice fed the HF/HS diet had lower body weight, increased insulin sensitivity, unchanged fasting glycemia, and reduced fasting insulinemia (**FIGURE 5B-E**) compared to CRE control mice. Again, female mice had profoundly reduced glucose-stimulated insulin secretion leading to modest glucose intolerance (**FIGURE 5F-G**). Conversely, male NCK1βKO mice fed the HF/HS diet tended to be heavier, while insulin sensitivity, fasting glycemia and fasting insulinemia remained unchanged compared to controls (**FIGURE 5H-K**). They had significantly increased glucose-stimulated insulin secretion accompanied by improved glucose tolerance (**FIGURE 5L-M**).

**Figure 5.**
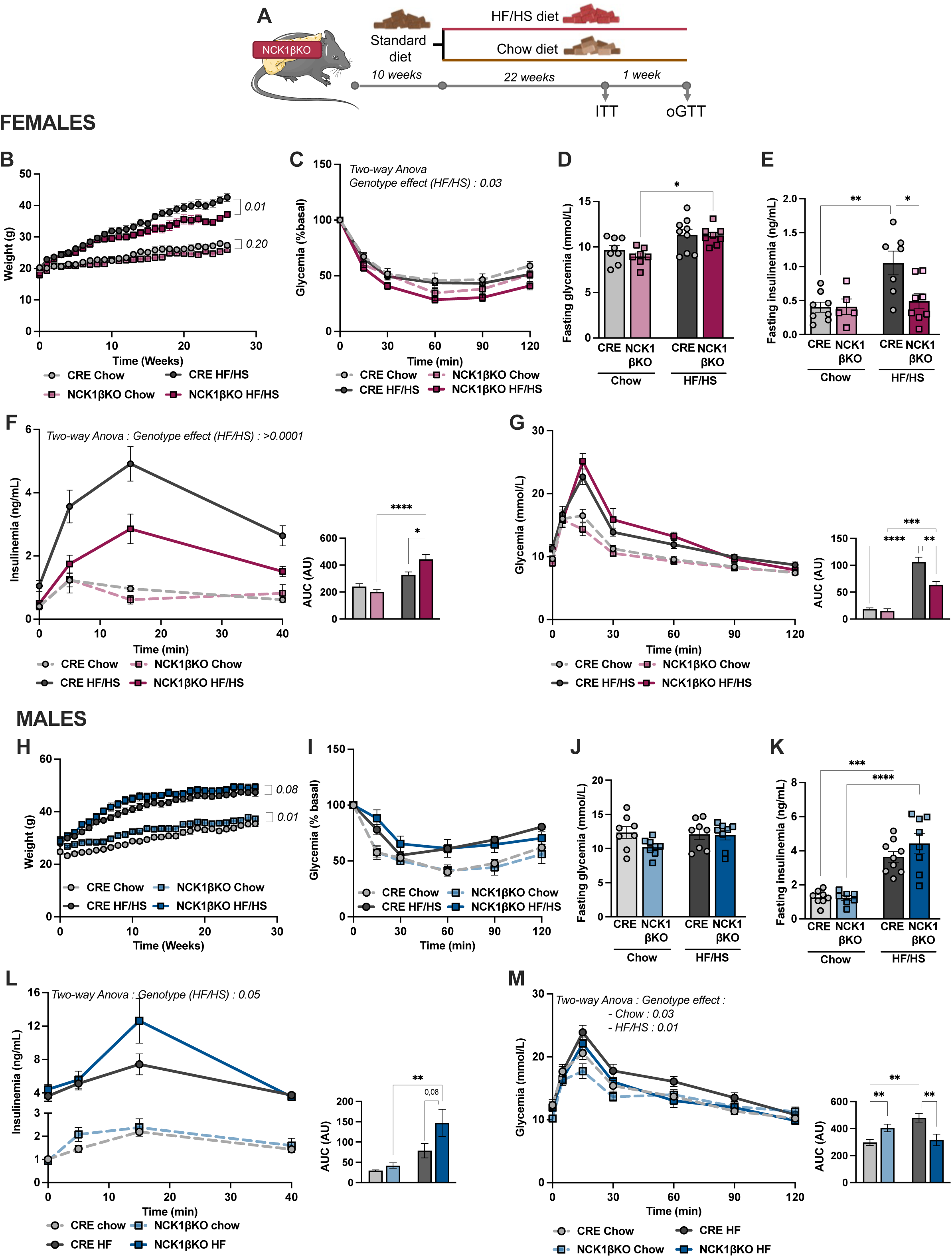
Long-term high-fat/high-sucrose feeding reveals sex-specific metabolic adaptation to β NCK1 loss. **(A**) Time course of the metabolic tests for female and male NCK1βKO and CRE control mice fed a chow or high fat high sucrose (HF/HS) diet. **(B, H)** Body weights of mice during the 22 weeks on diet (Females, n=7-9; Males, n= 8). **(C, I)** Insulin tolerance test, **(D, J)** fasting glycemia and **(E, K)** fasting insulinemia of 6 h-fast female and male NCK1βKO and CRE control mice fed a chow or HF/HS diet for 22 weeks (Females, n=5-9; Males, n=7-9). **(F, L)** Insulinemia and **(G, M)** glycemia during an oral glucose tolerance test of female and male NCK1βKO and CRE control mice after 23 weeks of chow or HF/HS diet (Females, n=5-9; Males, n= 7-8). Data are expressed as means ± SEM. Statistical significance was assessed using two-way ANOVA followed by Tukey’s multiple comparison with *p ≤ 0.05, **p ≤ 0.01, ***p ≤ 0.001 and ****p ≤ 0.0001.

Thus, our data from mice on either standard or HF/HS diets show that lack of β cell NCK1 causes differential disruption of serum insulin secretion between sexes. Consistent with the intrinsic GSIS defect, female NCK1βKO mice exhibited impaired insulin secretion that was exacerbated by metabolic stress and resulted in glucose intolerance. However, the unexpected *in vivo* increase in GSIS in male mice suggested a sex-dependent compensatory response that maintained, or even enhanced, insulin secretion and glucose homeostasis in response to β cell NCK1 loss.

### β cell NCK1 loss differentially impacts metabolic stress-induced changes in islet size and number in males and females

In mice, HF/HS diets increase insulin production and β cell mass as a compensatory response to insulin resistance and elevated metabolic load^4,29^. We therefore examined whether β cell NCK1 loss altered islet adaptation to metabolic stress in a sex-dependent manner. In line with differences *in vivo*, female NCK1βKO islets had reduced *Ins1/2* expression following exposure to glucolipotoxic conditions, while male NCK1βKO islets had increased expression (**FIGURE 6A-B**). Interestingly, both female and male NCK1βKO mice showed profound enhancement of pancreatic insulin content (**FIGURE 6C, G)** after 23 weeks of HF/HS feeding, but each showed a distinct islet remodeling response. Female NCK1βKO mice did not expand islet area compared to CRE control mice but had an increased proportion of small islets (**FIGURE 6D-F**). In contrast, male NCK1βKO mice exhibited marked increases in total islet area compared to CRE control mice driven by increased islet size (**FIGURE 6H-J**). Together, these findings suggest that NCK1 regulates sex-specific adaptive islet remodeling responses to metabolic stress.

**Figure 6.**
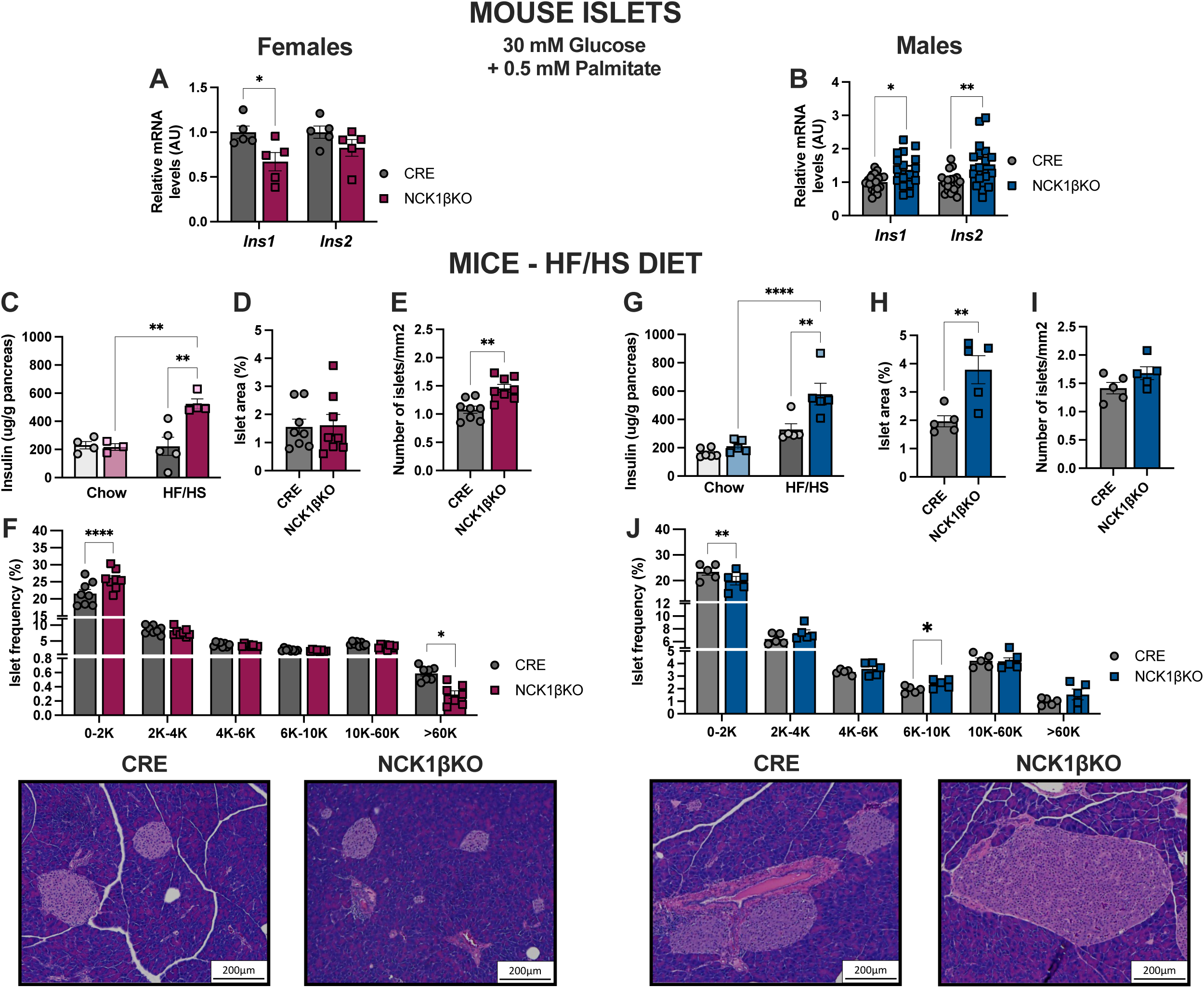
β cell NCK1 deficiency differentially affects islet size and density in male and female mice in response to metabolic stress. **(A, B)** Relative mRNA expression of insulin genes (*Ins1, Ins2*) in islets isolated from female and male NCK1βKO and CRE control mice, incubated 48 h in RPMI containing 30 mM glucose and 0,5 mM of palmitate (Females, n=5; Males, n=17-24). (**C, G**) Total pancreatic insulin content of female and male NCK1βKO and CRE control mice fed a chow or high-fat, high-sucrose (HF/HS) diet for 23 weeks (Females, n=3-5; Males, n=5-6). **(D, H)** % of islet area relative to total pancreatic area, **(E, I)** pancreatic islet density as the number of islets/mm^2^ of surface, **(F, J)** islet size distribution as percentage of pancreatic islets falling within each diameter range (μm), and representative images of H&E stained pancreas of female and male NCK1βKO and CRE control mice on a high-fat, high-sucrose (HF/HS) diet for 23 weeks (Females, n=8; Males, n=5). Data are expressed as means ± SEM. Statistical significance was assessed using two-way ANOVA followed by Tukey’s multiple comparison for panels A, C, G and unpaired t-test for others with *p ≤ 0.05, **p ≤ 0.01, ***p ≤ 0.001 and ****p ≤ 0.0001.

### NCK1 deficiency induces β cell proliferation in response to metabolic stress in a sex-dependent manner

The nearly two-fold increase in islet area observed only in male NCK1βKO mice suggested that NCK1 may regulate adaptive β cell mass expansion in a sex-dependent manner. HF/HS diet increases β cell mass mainly by enhancing proliferation^4,29,30^. Thus, we measured expression of key proliferation-associated genes in primary islets from NCK1βKO and CRE control mice exposed to high glucose or glucolipotoxic conditions. Female NCK1βKO islets showed no changes in proliferative gene expression in high glucose and an overall reduction in glucolipotoxic conditions, including 50% lower *CyclinA* and *CyclinB* mRNA. In contrast, male NCK1βKO islets exhibited modest, but significant increases in proliferative genes under both conditions **(FIGURE 7A-B)**. These data suggest that loss of NCK1 selectively enhances a cell-intrinsic, pro-proliferative transcriptional program in male islets. Taken together with differences in islet area and size, these data support a sex-dependent role for NCK1 in adaptive islet remodeling during metabolic stress.

**Figure 7.**
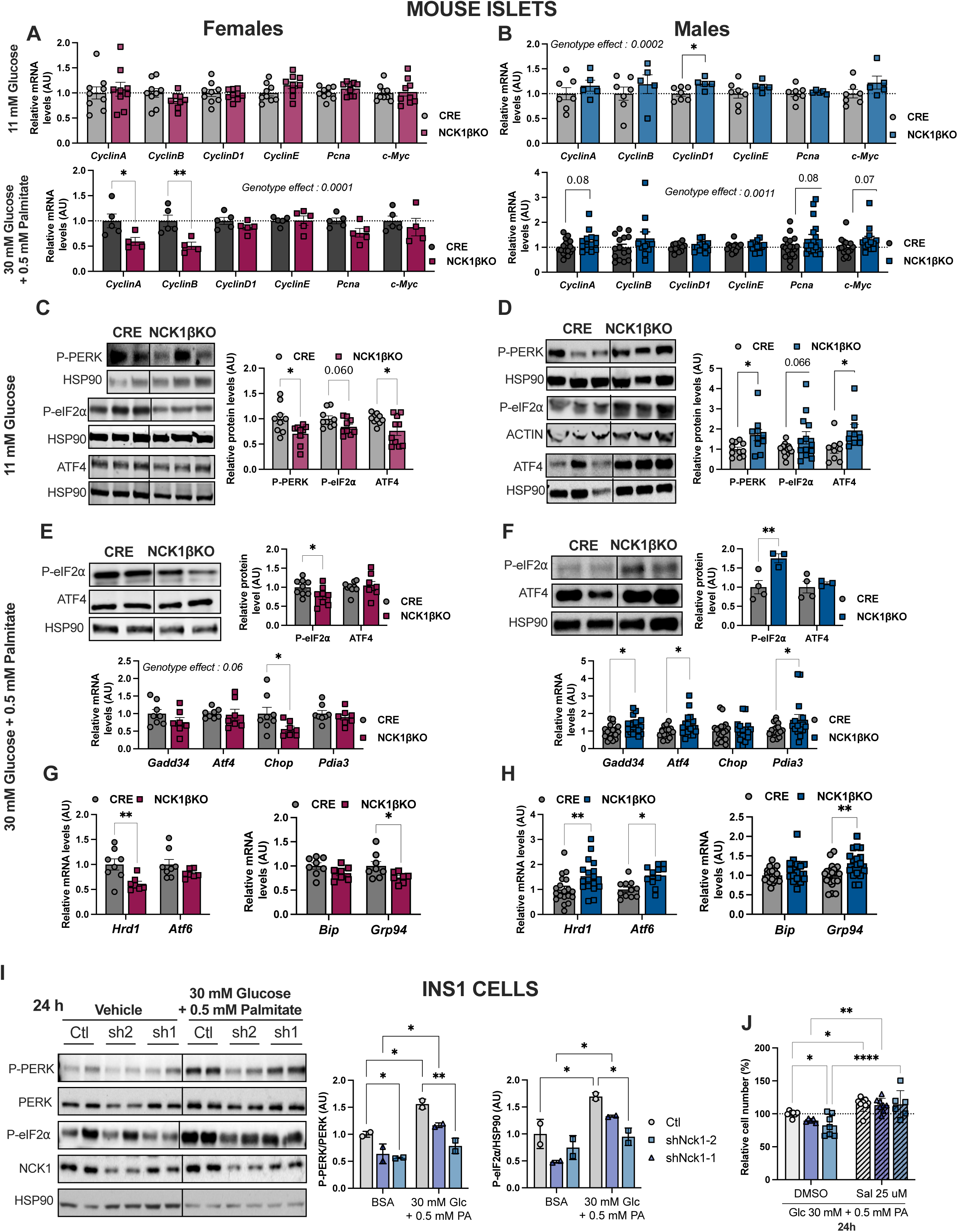
β cell NCK1 loss causes sex-specific changes in metabolic-stress induced cell proliferation and PERK signaling. **(A, B)** Relative mRNA expression of proliferative genes in islets isolated from female and male NCK1βKO and CRE control mice, incubated 48 h in RPMI containing 11 mM glucose (Females, n=9; Males, n=5-7) or 30 mM glucose + 0,5 mM palmitate (Females, n=4-5; Males, n=11-19). PERK pathway activation, measured by phosphorylation of PERK and/or eIF2*a*, ATF4 protein and/or relative mRNA expression of PERK pathway target genes in primary islets isolated from female and male NCK1βKO and CRE control mice, incubated 48 h in RPMI containing **(C, D)** 11 mM glucose (Females, n=9-10; Males, n=9-15) or **(E, F)** 30 mM glucose + 0,5 mM palmitate (Western blot: Females, n=8-9; Males, n=3-4 / qPCR: Females, n=7-8; Males, n=15-20). **(G, H)** IRE1 pathway targets *Hrd1* and *Atf6* and chaperone protein (*Bip* and *Grp94*) mRNA expression in islets isolated from female and male NCK1βKO and CRE control mice, incubated 48 h in RPMI with 30 mM glucose + 0,5 mM palmitate (Females, n=7-8; Males, n=7-19). **(I)** PERK pathway activation measured by phosphorylation of PERK and eIF2*a* in stable NCK1-KD or control INS-1 cells incubated 24 h in RPMI containing 11 mM glucose or 30 mM glucose + 0,5 mM palmitate (n=4). Western blot images and bar chart quantification are representative of 4 independent experiments. **(J)** Relative cell number, measured by MTT, of stable NCK1-KD or control INS-1 cells incubated 24 h in RPMI containing 11 mM glucose or 30 mM glucose + 0,5 mM palmitate with 25 μM salubrinal or vehicle control (DMSO). Data are percentage of viable cells relative to glucolipotoxic conditions of the same cell line, expressed as fold change of the control line (n=4). Data are expressed as means ± SD (INS-1) or SEM (islets). ACTIN or HSP90 were used as loading controls to normalize protein data, as specified. *Hprt* was used as an internal control to normalize qPCR. Statistical significance was assessed using multiple t-test analysis or two-way ANOVA followed by Tukey’s multiple comparison, with *p ≤ 0.05, **p ≤ 0.01, ***p ≤ 0.001 and ****p ≤ 0.0001.

### The PERK UPR response to glucolipotoxicity differs in male and female mice lacking β cell NCK1

We next explored mechanisms underlying sex-related differences in NCK1’s control of islet area and proliferative gene expression. Given that PERK signaling is a known regulator of β cell proliferation *in vivo*^28,31,32,33^, we examined activity of the PERK pathway in NCK1βKO male and female islets in response to metabolic stress. After 23 weeks of HF/HS feeding, females NCK1βKO islets maintained reduced PERK pathway activation compared to CRE control islets **(FIGURE 7C, E),** whereas male NCK1βKO islets now had increased PERK activity **(FIGURE 7D, F)**. A similar pattern was observed for additional UPR components, including *Hrd1*, *Atf6* and *Grp94* expression **(FIGURE 7G, H)**. This sex-specific regulation of the PERK pathway by NCK1 appeared specific to metabolic stress, as activation of PERK pathway by thapsigargin, which disrupts ER calcium homeostasis, was unaffected by NCK1 deficiency in INS-1 cells or primary islets (**FIGURE S5**).

Differential regulation of PERK by NCK1 in female and male islets during metabolic stress mirrored the sex-specific phenotypes in proliferative gene expression and islet area. We therefore asked whether effects of NCK1 loss on β cell proliferation could be through PERK. Under glucolipotoxic conditions, treatment of INS-1 cells with salubrinal (a PERK activator) increased both cell number and proliferation-associated gene expression, whereas a PERK inhibitor had the opposite effect (**FIGURE S6**). NCK1 deficiency under these conditions decreased the PERK pathway and cell number, whereas restoration of PERK activity with salubrinal rescued these defects (**FIGURE 7I,J**). Together, these findings support a role of NCK1 in regulating islet adaptation to metabolic stress through sex-specific regulation of PERK signaling, shaping islet remodeling and insulin secretion.

## DISCUSSION

In this study, we identify NCK1 as a regulator of β cell insulin production and secretion, proliferation and adaptation to metabolic stress, with PERK signaling emerging as a context-dependent downstream effector. NCK1 expression was induced by glucose but decreased in islets from people living with type 2 diabetes. Loss of NCK1 in β cells decreased insulin synthesis and glucose-stimulated insulin secretion *ex vivo* in islets from both sexes, which was associated with reduced PERK signaling. This was consistent with negative correlations between *NCK1*, *insulin* and PERK target gene expression in human β cells. NCK1 deletion triggered distinct sex-specific adaptive responses *in vivo,* exacerbated by metabolic stress. Female NCK1βKO mice exhibited the expected impairment of insulin secretion and glucose intolerance. This was accompanied with a persistent reduction of PERK signaling and increased proportion of small islets compared to control mice. Unexpectedly, male NCK1βKO mice had increased islet area, size and β cell proliferation, seemingly driven by enhancement of the PERK UPR branch, resulting in heightened *in vivo* insulin secretion and improved glucose tolerance. Together, these findings identify NCK1 as a sex-specific regulator of β cell adaptation to metabolic stress, potentially through differential modulation of PERK signaling and downstream islet remodeling responses.

Despite inducing similar intrinsic β cell defects in both sexes, NCK1 deficiency elicits distinct sex-specific adaptive strategies *in vivo*. Under basal conditions, these compensatory mechanisms were sufficient to preserve glucose homeostasis, likely because metabolic demand on β cells is limited within this context. Strikingly, HF/HS feeding exposed a clear divergence in their adaptative capacity to this stress. Female NCK1βKO mice appeared to rely primarily on increased insulin sensitivity and alternative islet remodeling towards increased proportion of small islets, which became inadequate to maintain glucose homeostasis as metabolic demand increased. In contrast, the male response, characterized by greater β cell expansion, was sufficient to overcome the intrinsic secretory defect and even enhanced *in vivo* insulin secretion and glucose control. These differential responses to NCK1 deficiency reflect sex-specific distinct adaptive strategies with different capacities to sustain glucose homeostasis under metabolic stress. This is consistent with growing evidence that β cell adaptation to metabolic stress is sexually dimorphic^34–37^ and suggests that NCK1 may play a role in these sex differences.

Our findings point to PERK signaling as one potential contributor to the β cell phenotypes associated with NCK1 loss, both under basal conditions and during metabolic stress. PERK, together with the adaptive UPR signaling, is required for multiple processes essential for β cell function and adaptation to increased metabolic demand, including insulin biosynthesis, β cell proliferation, differentiation and maintenance of β cell mass^4,7,28,31,32^. Under basal conditions, reduced PERK signaling in NCK1-deficient islets may contribute to the impaired insulin gene expression, insulin content and GSIS observed in both sexes. However, under metabolic stress, divergent regulation of PERK observed in male and female NCK1βKO mice could underlie their distinct adaptive responses. In males, enhanced PERK activation seems to promote an enhanced adaptive program, manifesting as increased insulin levels as well as proliferation-associated gene expression and islet expansion. Conversely, in females, persistently or permanently reduced PERK signaling may limit this response. Instead, female NCK1βKO mice had more small islets, suggesting a distinct adaptive remodeling program. Although the biological significance of this remodeling remains unclear, these small islets may arise from islet neogenesis^38^, representing newly formed or immature islets that require maturation before acquiring full glucose responsiveness^39–41^. Thus, these small islets would contribute to the increased pancreatic insulin stores observed in female NCK1βKO mice, but may remain functionally immature and unable to restore insulin secretion. Reduced PERK signaling may also contribute to this phenotype, as PERK deficiency has been associated with reduced islet size and impaired β cell maturation^42,43^. Overall, these findings support a potential role for PERK in the sex-specific phenotype and adaptive responses to β cell NCK1 loss, although its causal contribution remains to be established.

A link between NCK1 and PERK has been previously established. However, our observations contrast with previous studies showing β cell NCK1 deficiency enhances basal PERK signaling and proinsulin content in MIN6 cells^8,9^. These discrepancies may arise from differences in experimental models and metabolic environment, which is consistent with previous studies indicating that the relationship between NCK1 and PERK can vary depending on the cellular context^9,18,19,33^. Our findings extend this concept and suggest that metabolic and hormonal context may also influence this relationship. Interestingly, estrogen signaling promotes β cell function and survival, while enhancing adaptive UPR signaling and resilience to ER stress^44–46^. Our data raise the possibility that NCK1 lies downstream of estrogen signaling in these roles. Together, these findings raise the possibility that NCK1 may act as a context-dependent modulator of the UPR, integrating metabolic and hormonal cues to influence β cell adaptation, although the mechanisms underlying its sex-specific regulation of PERK remain to be established.

Our findings showing the glucose responsiveness of NCK1 expression, together with its reported role in glucose metabolism, such as promoting glucose uptake, glycolysis and oxidative phosphorylation in primary human T cells^47^, suggests that NCK1 may also contribute to glucose sensing in β cells. Thus, beyond PERK signaling, NCK1 may play a role in either glucose sensing or metabolism in β cells, supporting the glucose-coupled-insulin secretory machinery. NCK1 activates PI3K/AKT^23,48^ and ERK signaling^49,50^, both of which positively regulate insulin gene expression in β cells^51,52^. NCK1 also regulates protein translation initiation^20,53^ and actin cytoskeleton remodeling^15,16,18^, processes that are essential for hormone biosynthesis and granule exocytosis^58,59^. Given its scaffolding function of NCK1, it likely coordinates multiple signaling and cellular processes to maintain β cell functional capacity and adaptation. How these pathways are regulated and interconnected remains to be determined.

Finally, reduced β cell NCK1 in type 2 diabetes raises the possibility that this may contribute to β cell dysfunction by impairing insulin production and secretion, as well as altering adaptation to metabolic stress. This may be particularly relevant during diabetes progression, when β cells must increase insulin production and adapt their functional mass to compensate for insulin resistance. Importantly, we see NCK1 decreased in both male and female donors, but it remains to be tested whether the sex-differences seen in mice translate to humans. Determining whether reduced NCK1 contributes directly to human β cell failure, and whether restoring NCK1 activity can improve their function or survival, will be important future directions and could reveal new strategies to preserve insulin secretion and β cell mass in diabetes.

## Conclusion

In conclusion, our findings identify NCK1 as an important regulator of β cell functional capacity and adaptation to metabolic stress. Beyond its role in maintaining insulin production and secretion, NCK1 appears to shape sex-specific adaptive responses that determine the ability of β cells to preserve glucose homeostasis under increased metabolic demand. Impaired NCK1-dependent adaptation may contribute to β cell dysfunction during diabetes progression. More broadly, our study highlights NCK1 as a potential link between β cell stress signaling, remodeling and the sex-dependent regulation of β cell adaptation, important aspects of diabetes pathogenesis.

## ACKNOWLEDGEMENTS

We thank members of the IRCM animal, microscopy, and molecular biology core facilities for invaluable technical assistance.

## DATA AVAILABILITY

The datasets generated and/or analyzed during the current study are available from the corresponding author upon a request. Human β cell single-cell RNA-sequencing data analyzed in this study are publicly available through the Gene Expression Omnibus (GEO) under accession number GSE221156 and through Zenodo.

## AUTHOR CONTRIBUTIONS

LM, NJ and JLE designed concept and experiments. LM performed experiments and analyzed the data. RH, JT, LG, CB, MG and CS assisted with experimentation. MS performed the bioinformatics analysis of the scRNAseq data. LM and JLE wrote the manuscript. CS and AG contributed to study conception, provided intellectual input and critically revised the manuscript. All authors reviewed the manuscript.

## ABBREVIATIONS

ATF4: activating transcription factor 4
ATF6: activating transcription factor 6
BIP: binding immunoglobulin protein
βKO: β cell-specific knockout
CRE: Ins1-Cre control mice
ER: endoplasmic reticulum
GSIS: glucose-stimulated insulin secretion
HF/HS: high-fat/high-sucrose
HRD1: HMG-CoA reductase degradation protein 1
IRE1α: inositol-requiring enzyme 1α
MMTT: mixed meal tolerance test
NCK1: non-catalytic region of tyrosine kinase adaptor kinase 1
oGTT: oral glucose tolerance test
PERK: protein kinase RNA-like endoplasmic reticulum kinase (EIF2AK3)
P-eIF2α: phosphorylated eukaryotic initiation factor 2α
scRNA-seq: single-cell RNA sequencing
SEM: standard error of the mean
T2D: type 2 diabetes
UPR: unfolded protein response

