## Supplementary Figures and Tables for "β-cell NCK1 is reduced in type 2 diabetes, impairing the unfolded protein response and insulin secretion while revealing sex-specific adaptation"

<u>**SUPPLEMENTARY FIGURES & LEGENDS**</u>

### FIGURE S1

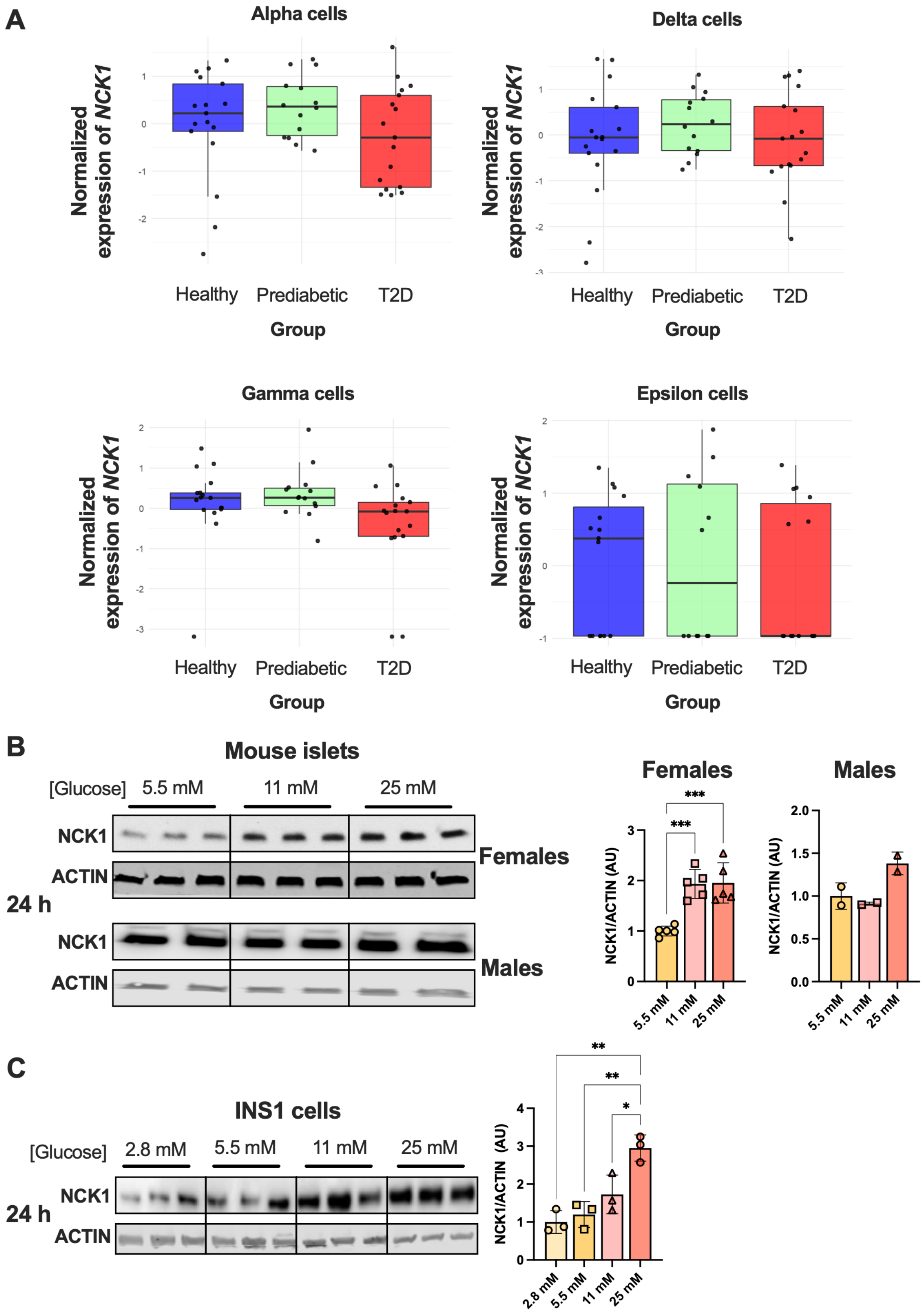

A
Alpha cells
Delta cells
Gamma cells
Epsilon cells
Normalized expression of NCK1
Healthy
Prediabetic
T2D
Group
B
Mouse islets
[Glucose]
5.5 mM
11 mM
25 mM
NCK1
ACTIN
24 h
Females
Males
NCK1/ACTIN (AU)
C
INS1 cells
2.8 mM

**Figure S1. NCK1 expression is unaffected by T2D status in human donor islet non-β cells and is induced by glucose in mouse islets.**

**(A)** Pseudobulk analysis of *NCK1* mRNA expression in human α, δ, γ and ε endocrine cell populations from healthy (n=17), prediabetic (n=14) and type 2 diabetic (n=17) donors. **(B)** Relative protein expression of NCK1 in female and male mouse islets incubated for 48 h with increasing glucose concentrations (5.5, 11 and 25 mmol/L) following overnight recovery in RPMI containing 5.5 mmol/L glucose. Representative western blots and corresponding quantification are shown. **(C)** Relative protein expression of NCK1 in INS-1 cells incubated for 24 h with increasing glucose concentrations (2.8, 5.5, 11 and 25 mmol/L). Representative western blots and quantification are shown. ACTIN was used as loading control. Data are expressed as mean ± SD or SEM as indicated. Statistical significance was assessed using one-way ANOVA with Tukey's multiple comparisons test. $*p \leq 0.05$, $**p \leq 0.01$, $***p \leq 0.001$.

FIGURE S2

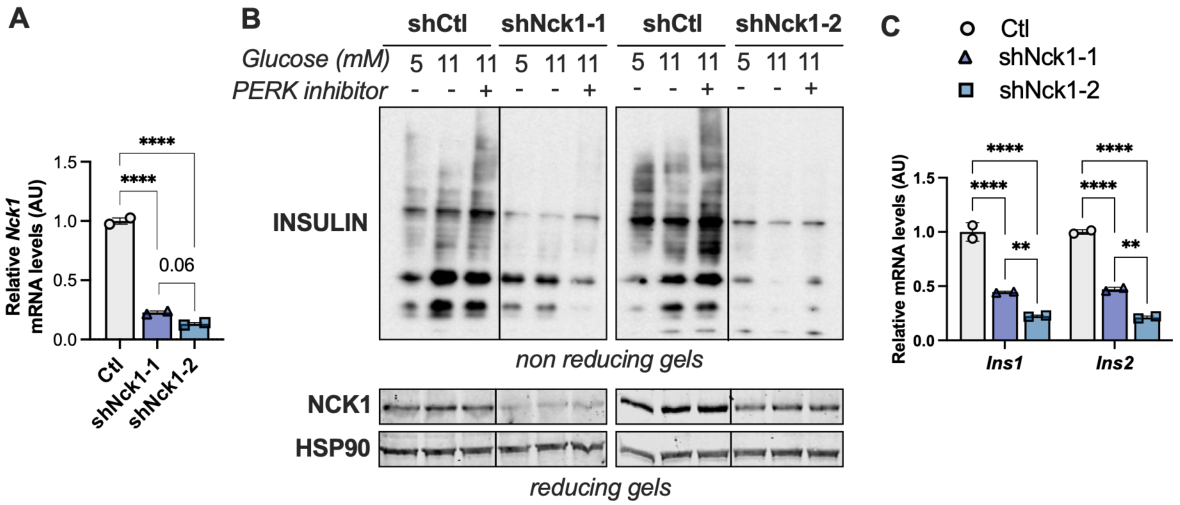


**Figure S2. Acute NCK1 silencing reduces insulin expression and proinsulin/insulin content in INS-1 cells.**

**(A)** Relative *Nck1* mRNA expression in INS-1 cells transduced with control shRNA (Ctl) or two independent shRNAs targeting *Nck1*(shNck1-1 and shNck1-2). **(B)** Relative expression of insulin genes (*Ins1* and *Ins2*) in control and NCK1-deficient INS-1 cells. **(C)** Representative immunoblots of insulin, proinsulin folding assessed by reducing and non-reducing SDS-PAGE, and NCK1 protein expression in control and NCK1-deficient INS-1 cells incubated in 5 or 11 mmol/L glucose with or without PERK inhibition (as a control). Data are expressed as mean ± SD. Statistical analyses were performed using one-way or two-way ANOVA as appropriate. $^{**}p \leq 0.01$, $^{***}p \leq 0.001$, $^{****}p \leq 0.0001$.

FIGURE S3

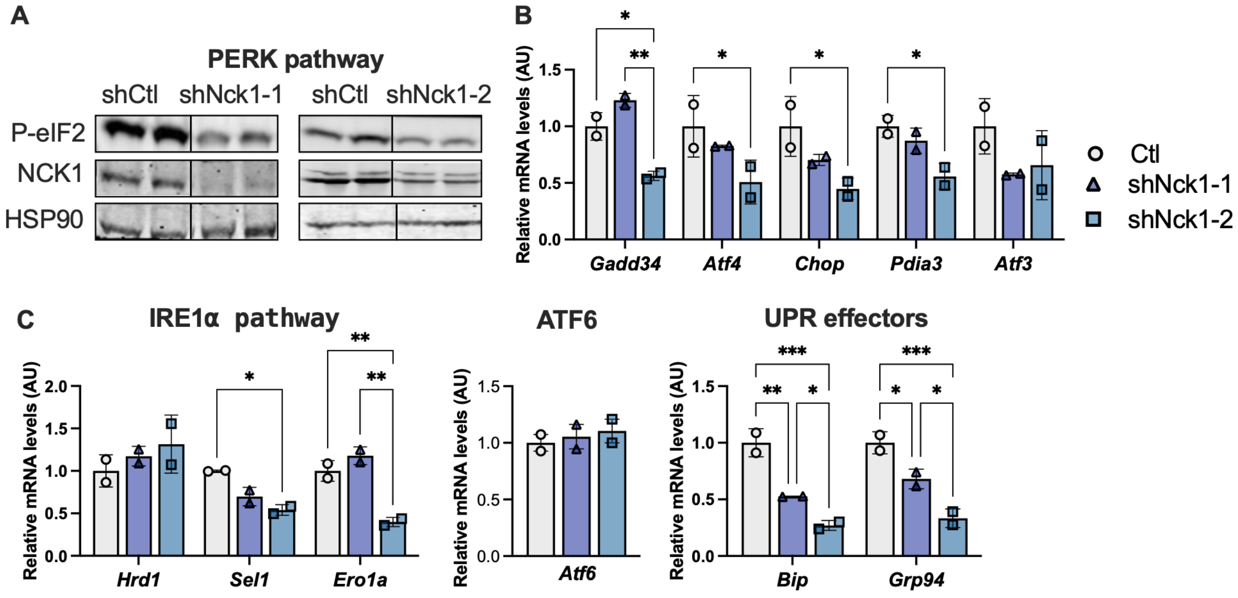


**Figure S3. NCK1 deficiency alters UPR signaling in INS-1 cells.**

**(A)** Relative mRNA expression of PERK pathway target genes (*Gadd34, Atf4, Chop, Pdia3,* and *Atf3*) in control and stable NCK1-KD INS-1 cells. **(B)** Relative expression of genes associated with the IRE1α pathway (*Hrd1, Sel1,* and *Ero1a*). **(C)** Relative mRNA expression of *Atf6* and ER chaperones (*Bip* and *Grp94*). Representative immunoblot showing phosphorylated eIF2α and NCK1 protein expression is presented. Data are expressed as mean ± SD. Statistical significance was assessed using one-way ANOVA with Tukey's multiple comparisons test. $^*p \leq 0.05$, $^{**}p \leq 0.01$, $^{***}p \leq 0.001$.

FIGURE S4

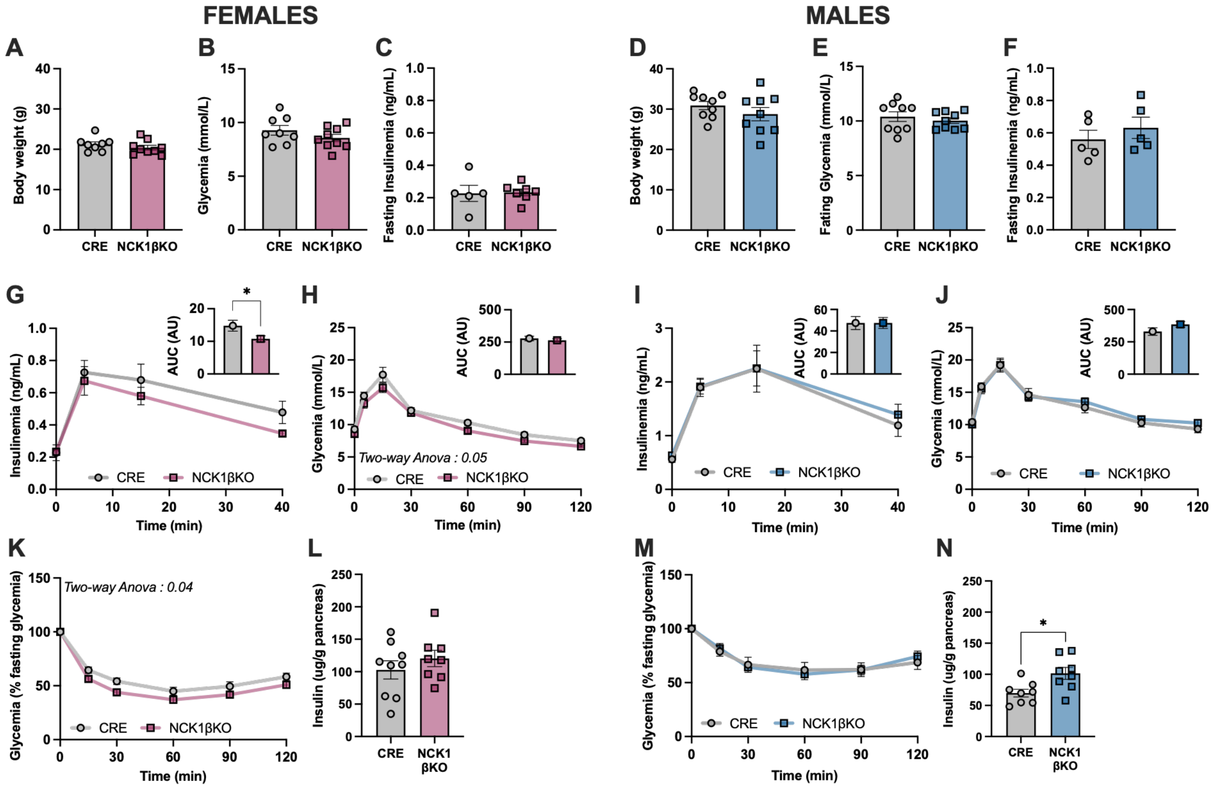


**Figure S4. β-cell NCK1 deletion has little effect on glucose homeostasis under chow-fed conditions.**

**(A, B)** Body weight of female and male NCK1βKO and CRE control mice maintained on a chow diet. **(C, D)** Fasting glycemia and fasting insulinemia in female mice. **(E, F)** Fasting glycemia and fasting insulinemia in male mice. **(G, I)** Serum insulin levels during oral glucose tolerance tests (oGTT) in female and male mice, respectively. **(H, J)** Blood glucose excursions and corresponding area under the curve (AUC) during oGTT in female and male mice. **(K, M)** Insulin tolerance tests performed in female and male mice. **(L, N)** Total pancreatic insulin content in female and male mice. Data are expressed as mean ± SEM. Statistical significance was assessed using unpaired *t*-tests or two-way ANOVA followed by Tukey's multiple comparisons test. *$p \leq 0.05$.

FIGURE S5

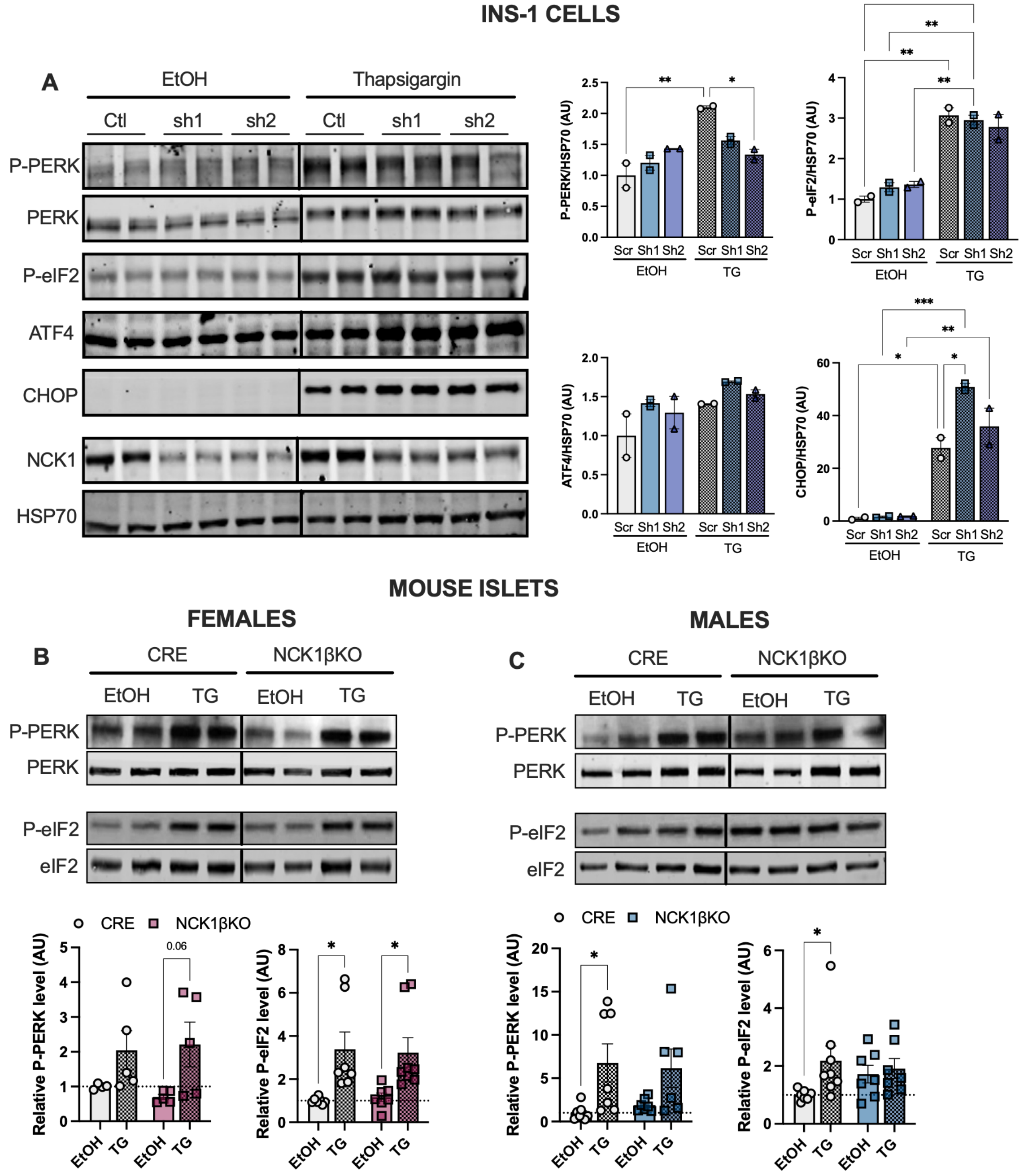


**Figure S5. NCK1 deficiency does not impair thapsigargin-induced PERK activation.**

**(A)** Representative immunoblots and quantification of PERK pathway activation measured by phosphorylated PERK and phosphorylated eIF2α in control or NCK1-deficient INS-1 cells treated with vehicle (EtOH) or thapsigargin (TG). ATF4 and CHOP protein levels are also shown. **(B, C)** Representative immunoblots and quantification of PERK activation in female and male islets isolated from CRE control and NCK1βKO mice treated with vehicle or thapsigargin. Data are expressed as mean ± SD (INS-1 cells) or SEM (islets). Statistical significance was assessed using two-way ANOVA followed by Tukey's multiple comparisons test. $*p \leq 0.05$, $**p \leq 0.01$.

FIGURE S6

INS-1 CELLS

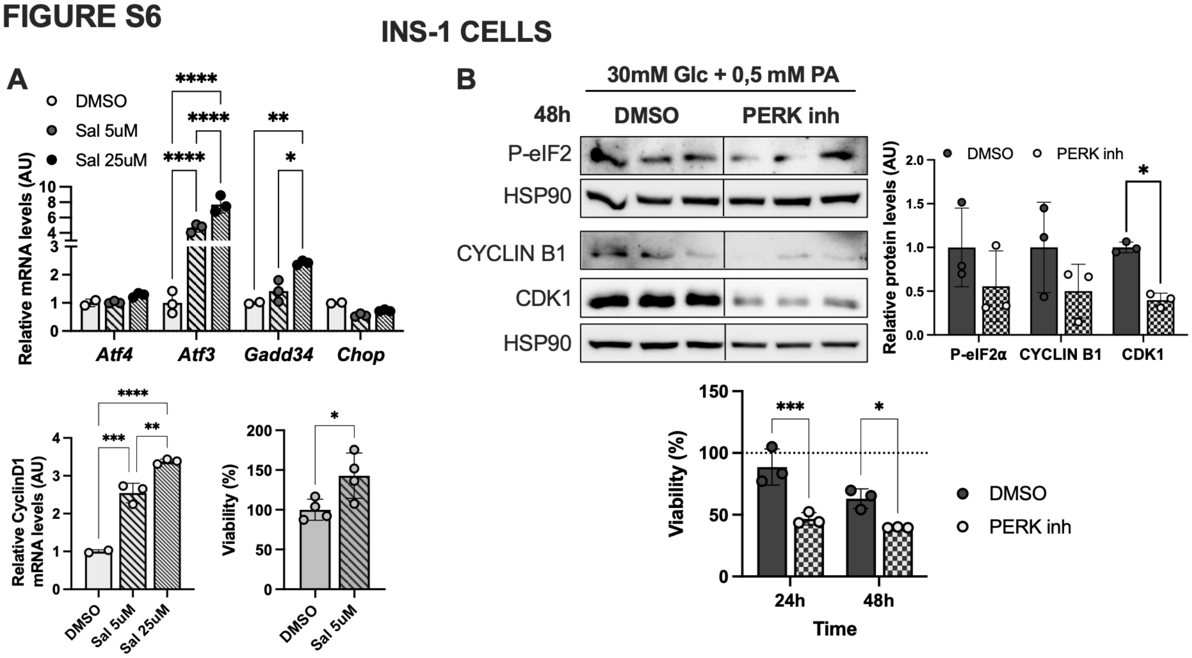


**Figure S6. Pharmacological modulation of PERK signaling regulates β-cell proliferation and survival under glucolipotoxic stress.**

**(A)** Relative expression of PERK target genes (*Atf4, Atf3, Gadd34,* and *Chop*) in INS-1 cells treated with vehicle (DMSO) or salubrinal (5 or 25 µM). Relative *Cyclin D1* mRNA expression after PERK activation with salubrinal. Cell viability measured by MTT assay in INS-1 cells exposed to glucolipotoxic conditions and treated with salubrinal or PERK inhibitor. **(B)** Representative immunoblots and quantification of P-eIF2α, Cyclin B1, and CDK1 protein levels following PERK inhibition. Data are expressed as mean ± SD. Statistical significance was determined using one-way or two-way ANOVA with Tukey's multiple comparisons test. *p ≤ 0.05, **p ≤ 0.01, ***p ≤ 0.001, ****p ≤ 0.0001.

### SUPPLEMENTARY TABLES

**Supplementary Table 1: shRNA constructs used for NCK1 knockdown.**

| **shRNAs** | **Sequences** |
|---|---|
| Sh-Ctl | 5'-GGTTCAGATGTGCGGCGAGT-3' |
| Sh-Nck1-1 | 5'-GCAGTTGTCAATAACCTAAAT-3' |
| Sh-Nck1-2 | 5'-GCTGGCAATCCTTGGTATTAT-3' |

**Supplementary Table 2: Antibodies and experimental conditions.**

| Antibody | Origin | Reference | Host | Dilution |
|---|---|---|---|---|
| Beta – ACTIN | Sigma | A5441 | Mouse | 1/5000 |
| ATF4 | Proteintech | 10835-1-AP | Rabbit | 1/1000 |
| CHOP | Cell signaling | 2895 | Mouse | 1/1000 |
| eIF2a | Cell signaling | 2103 | Mouse | 1/1000 |
| P-eIF2a (S51) | Cell signaling | 9721 | Rabbit | 1/1000 |
| CYCLIN B1 | Santa Cruz Biotechnology | sc-245 | Mouse | 1:1000 |
| CDC2 p34 (CDK1) | Santa Cruz Biotechnology | sc-54 | | 1/500 |
| HSP90 | Cell signaling | 4874 | Rabbit | 1/1000 |
| HSP70 | Cell signaling | 4872 | Rabbit | 1/1000 |
| INSULIN | Cell signaling | 8138 | Mouse | 1/1000 |
| NCK1 | Gift from Dr. Louise Larose | Latreille *et al.*, 2011 | Rabbit | 1/1000 |
| P-PERK (Thr980) | Cell signaling | 3179 | Rabbit | 1/500 |
| PERK | Cell signaling | 3192 | Rabbit | 1/1000 |
| Goat anti-mouse HRP | BioRad | 1706516 | Goat | 1:5000 |
| Goat anti-rabbit HRP | BioRad | 1706515 | Goat | 1:5000 |
| Goat anti-mouse IRDye800CW | Li-cor | 925-32210 | Goat | 1:15000 |
| Goat anti-rabbit IRDye800CW | Li-cor | 925-32211 | Goat | 1:15000 |
| Goat anti-mouse IRDye680CW | Li-cor | 926-68070 | Goat | 1:15000 |
| Goat anti-rabbit IRDye680CW | Li-cor | 926-68071 | Goat | 1:15000 |

**Supplementary Table 3: Primers used for quantitative real-time PCR.**

| Genes | Fwd | Rev | Species |
|---|---|---|---|
| *Atf3* | CTGCTGCCAAGTGTCGAAACA AGA | AGTTCGGCATTCACACTCTCC AGT | Rat |
| *Atf4* | GCCAAGCACTTCAAACCTCA | CAATCTGTCCCGGAAAAGGC | Rat |
| *Atf4* | GTCCCTTTCCTCTTCCCCTC | GATTTCGTGAAGAGCGCCAT | Mouse |
| *Atf6* | CCGGTTCTTCCTCATGGACC | GGTCTGACTCCCAAGGCATC | Mouse & rat |
| *Bip* | TCGACTTGGGGACCACCTAT | AGTGAAGGCCACATACACG | Mouse & rat |
| *Chop* | GCAGCGACAGAGCCAGAATA | ATGTGCGTGTGACCTCTGTT | Mouse |
| *Chop* | AGTCTCTGCCTTTCGCCTTTGA GA | TGCAGGGTCAAGAGTAGTGAA GGT | Rat |
| *Cyclin A* | CACTGACACCTCTTGACTATCC | CGTTCACTGGCTTGTCTTCTA | Mouse |
| *Cyclin B* | GGTCGTGAAGTGACTGGAAA | GTCTCCTGAAGCAGCCTAAAT | Mouse |
| *Cyclin D1* | CAGAGGCGGATGAGAACAAG | GAGGGTGGGTTGGAAATGAA | Mouse & rat |
| *Cyclin E* | GGTTATCAGTGGTGCGACATAG | GGAAGTGCTTGAGCTTGGA | Mouse |
| *Edem* | CTACCTGCGAAGAGGCCG | GTTCATGAGCTGCCCACTGA | Mouse & rat |
| *Ero1-a* | CTTATATCTGGCCTGCACGC | TCTGTGACATTGTGACCCCA | Mouse & rat |
| *Gadd 34* | GAGAAGACCAAGGGACGTGG | TCGATCTCGTGCAAACTGCT | Mouse & rat |

| | | | |
|---|---|---|---|
| *Grp94* | CACTCAAATCGAACACGGCTTGCT | AGAAGATTCCGCCTCCTTTCTGCT | Mouse & rat |
| *Herpud1* | TCCCAAAGACGCCAAGTGTC | TTGGGACTCTTCACCAGCAG | Mouse & rat |
| *Hprt* | CCAACAGAGGGCCACAATGT | TGAAAGACTTGCTCGAGATGTCA | Mouse & rat |
| *Hrd1* | TTTATGGAACGCAGCCCCAA | AAGCCTGTAAACAGCTCCGT | Mouse & rat |
| *Ins1* | GAAGTGGAGGACCCACAAGTG | ATCCACAATGCCACGCTTCT | Mouse & rat |
| *Ins2* | GAAGTGGAGGACCCACAAGTG | GATCTACAATGCCACGCTTCTG | Mouse & rat |
| *Mafa* | AGGAGGAGGTCATCCGACTG | CTTCTCGCTCTCCAGAATGTG | Mouse |
| *c-Myc* | CTCCGTACAGCCCTATTTCATC | TGGGAAGCAGCTCGAATTT | Mouse |
| *Pcna* | TCCTGTGCAAAGAATGGGGT | TAGGAGACAGTGGAGTGGCTT | Mouse |
| *Pdia3* | AGCCAATGATGTGCCTTCTC | TTTAATTCACGGCCACCTTC | Mouse & rat |
| *Pdx1* | CTTAACCTAGGCGTCGCACAA | GAAGCTCAGGGCTGTTTTTCC | Mouse |
| *Sel1* | GAAGATGGCAGACTGTGGTGT | GCATCTGTCGTCTTTTGGCAG | Mouse & rat |

### Supplementary Table 4: Human Islet Preparations Used in Research Study

| **Islet preparation** | **1** | **2** | **3** | **4** | **5** |
|---|---|---|---|---|---|
| Unique identifier | R186 (SAMN41425537) | R253 (SAMN41425604) | R159 (SAMN41425510) | R195 (SAMN41425546) | R139 (SAMN41425490) |
| Donor age (years) | 67 | 57 | 60 | 54 | 50 |
| Donor sex (M/F) | M | M | M | M | M |
| Donor BMI ($kg/m^2$) | 28,1 | 25,6 | 30,4 | 35 | 24.8 |
| Donor $HbA_{1c}$ | 5.6 | 5 | 5.5 | 5.5 | 5.9 |
| Origin/source of islets[b] | Alberta IsletCore | Alberta IsletCore | Alberta IsletCore | Alberta IsletCore | Alberta IsletCore |
| Islet isolation centre | Alberta Diabetes Institute | Alberta Diabetes Institute | Alberta Diabetes Institute | Alberta Diabetes Institute | Alberta Diabetes Institute |
| Donor history of diabetes? | No | No | No | No | No |
| Diabetes duration (years) | | | | | |
| Glucose-lowering therapy | | | | | |
| Donor cause of death | CVA/stroke | | | Anoxia | |
| Cold ischaemia time (h) | 15h | 14h | 2h | 17h | 16h |
| Estimated purity (%) | 95% | 90% | 70% | 95% | 90% |
| Handpicked to purity? | Yes | Yes | Yes | Yes | Yes |

| 6 | 7 | 8 | 9 | 10 | 11 |
|---|---|---|---|---|---|
| R125 (SAMN41425476) | R131 (SAMN41425482) | R213 (SAMN41425564) | R170 (SAMN41425521) | R236 (SAMN41425587) | R263 (SAMN41425614) |
| 70 | 52 | 57 | 49 | 51 | 71 |
| M | M | M | M | M | M |
| 29.4 | 26 | 23.1 | 38.6 | 35.3 | 38.3 |
| 7.7 | 6.8 | 6.7 | 6.5 | 8.6 | 7.6 |
| Alberta IsletCore | Alberta IsletCore | Alberta IsletCore | Alberta IsletCore | Alberta IsletCore | Alberta IsletCore |
| Alberta Diabetes Institute | Alberta Diabetes Institute | Alberta Diabetes Institute | Alberta Diabetes Institute | Alberta Diabetes Institute | Alberta Diabetes Institute |
| Yes | Yes | Yes | Yes | Yes | Yes |
| 20 | 2 | 22 | 3 | Non compliant T2D, DTZ negative | 9 |
| Metformin | Metformin | Metformin | | | |
| stroke | Intracranial bleed | | Cardiac arrest | | |
| 14h | 8.6h | 16h | 11h | 23h | 20h |
| 75% | 70% | 50% | 85% | | 75% |
| Yes | Yes | Yes | Yes | Yes | Yes |

| **Islet preparation** | **12** | **13** | **14** | **15** | **16** |
|---|---|---|---|---|---|
| Unique identifier | R246 (SAMN41425597) | R190 (SAMN41425541) | R234 (SAMN41425585) | R164 (SAMN41425515) | R191 (SAMN41425542) |
| Donor age (years) | 65 | 66 | 50 | 44 | 54 |
| Donor sex (M/F) | F | F | F | F | F |
| Donor BMI ($kg/m^2$) | 39.2 | 33.1 | 31.7 | 30.8 | 30.6 |
| Donor $HbA_{1c}$ | 5.8 | 6.2 | 5.7 | 6 | 6.6 |
| Origin/source of islets | Alberta IsletCore | Alberta IsletCore | Alberta IsletCore | Alberta IsletCore | Alberta IsletCore |
| Islet isolation centre | Alberta Diabetes Institute | Alberta Diabetes Institute | Alberta Diabetes Institute | Alberta Diabetes Institute | Alberta Diabetes Institute |
| Donor history of diabetes? | No | No | No | No | Yes |
| Diabetes duration (years) | | | | | 10 |
| Donor cause of death | Spontaneous intracerebral hemorrhage | Stroke | Anoxia | | Anoxia |
| Cold ischaemia time (h) | 19h | 20h | 16h | 9.3h | 14h |
| Estimated purity (%) | 75% | 95% | 90% | 85% | 40% |
| Handpicked to purity? | Yes | Yes | Yes | Yes | Yes |

| Islet preparation | 17 | 18 | 19 | 20 |
|---|---|---|---|---|
| Unique identifier | R240 (SAMN41425591) | R244 (SAMN41425595) | R231 (SAMN41425582) | R137 (SAMN41425488) |
| Donor age (years) | 55 | 48 | 41 | 56 |
| Donor sex (M/F) | F | F | F | F |
| Donor BMI (kg/m$^2$) | 30.9 | 30.4 | 37.1 | 36.7 |
| Donor $HbA_{1c}$ | 6.9 | 7.5 | 6.8 | 5.5 |
| Origin/source of islets | Alberta IsletCore | Alberta IsletCore | Alberta IsletCore | Alberta IsletCore |
| Islet isolation centre | Alberta Diabetes Institute | Alberta Diabetes Institute | Alberta Diabetes Institute | Alberta Diabetes Institute |
| Donor history of diabetes? | Yes | Yes | Yes | Yes |
| Diabetes duration (years) | Prediabetic | 2 | 0.5 | 2.5 |
| Glucose-lowering therapy at time of death | On medical chart | Metformin | Diet only | Metformin |
| Donor cause of death | | | Aneurysmal rupture | Intracranial bleed |
| Cold ischaemia time (h) | 13h | 12h | 18h | 2h |
| Estimated purity (%) | 50% | 70% | 50% | 90% |
| Handpicked to purity? | Yes | Yes | Yes | Yes |
